# Evolution of T cell responses in the tuberculin skin test reveals generalisable Mtb-reactive T cell metaclones

**DOI:** 10.1101/2025.04.12.648537

**Authors:** Carolin T Turner, Andreas Tiffeau-Mayer, Joshua Rosenheim, Aneesh Chandran, Rishika Saxena, Ping Zhang, Jana Jiang, Michelle Berkeley, Flora Pang, Imran Uddin, Gayathri Nageswaran, Suzanne Byrne, Akshay Karthikeyan, Werner Smidt, Paul Ogongo, Rachel Byng-Maddick, Santino Capocci, Marc Lipman, Heinke Kunst, Stefan Lozewicz, Veron Rasmussen, Gabriele Pollara, Julian C Knight, Alasdair Leslie, Benjamin M Chain, Mahdad Noursadeghi

## Abstract

T cells contribute to immune protection and pathogenesis in tuberculosis, but measurements of polyclonal responses have failed to resolve correlates of outcome. We report the first temporal evaluation of the human *in vivo* clonal repertoire of Mtb-reactive T cell responses, by T cell receptor (TCR) sequencing at the site of a standardised antigenic challenge. Initial recruitment of non-Mtb reactive T cells is followed by enrichment of Mtb-reactive clones arising from oligoclonal T cell proliferation. We introduce a modular computational pipeline, Metaclonotypist, to sensitively cluster distinct TCRs with shared epitope specificity, which we apply here to establish a catalogue of public Mtb-reactive HLA-restricted T cell metaclones. Although most *in vivo* Mtb-reactive T cells are private, 10 metaclones were sufficient to identify Mtb-T cell reactivity across our study population (N≥128), indicating striking population level immunodominance of specific TCR-peptide interactions that may offer novel approaches to patient stratification and vaccine development.

## Background

*Mycobacterium tuberculosis* (Mtb) remains the commonest microbial cause of death worldwide, but most incident infections do not progress to tuberculosis (TB) disease^1,2^. Immunodeficiency is associated with increased risk of disease, indicating a role for protective immunity. However, disease is mediated by immunopathology, associated with a failure to restrict Mtb growth. Better understanding of the immune correlates of protection and pathogenesis remain global research priorities to inform novel approaches to disease-risk stratification in Mtb infected people, vaccine development and evaluation, and identification of targets for host-directed therapies.

T cells are essential for protective immunity to TB. They are thought to augment bacterial restriction within intracellular niches such as macrophages^3,4^. T cell-mediated protection against TB is evident in increased disease risk associated with genetic deficiencies of IL-12 and IFNγ signalling^5^, T cell depletion in people living with HIV^6^, and experimental T cell depletion in non-human primates^7,8^. Yet, frequency of circulating Mtb-reactive T cells, and limited analysis of their functional attributes (cytokine production or cytolytic degranulation) do not predict natural or vaccine-inducible protective immunity in humans^4^. We have also reported evidence for T cell-mediated pathogenesis in TB, illustrated by enrichment of IL-17 producing T cells in people with disease compared to those who have controlled infection^9^, disease triggered by checkpoint inhibitor therapies that increase effector T cell function^10^ and direct stimulation of Mtb-growth by IFNγ^11^.

T cells exist as clonal populations identified by their T cell receptor (TCR), most commonly composed of αβ heterodimers produced by imprecise somatic gene recombination during T cell development and responsible for signalling T cell activation following recognition of antigen bound to MHC molecules. Mtb proteome-wide studies have identified immunodominant protein antigens^12–14^. To date, use of whole protein or pooled peptide antigens to quantify Mtb-reactive T cells has not resolved correlates of protection and pathogenesis, potentially because they measure polyclonal responses in which responses to distinct peptide-MHC targets have differential effects on outcome. TCR sequencing enables an antigen-agnostic approach to resolve clonal T cell responses. This has provided proof of concept for potential correlates of outcome^15^, but the generalisability of these findings is not known.

Importantly, studies of human T cell biology in TB have relied heavily on investigation of Mtb reactive T cells from blood samples that are limited by sampling depth, because they contain <0.001% of the T cell clonal repertoire of an individual^16^, only a small fraction of which is Mtb reactive. Alternatively, investigation of T cells from the site of disease, such as bronchoalveolar lavage specimens or tissue biopsies, can enrich for the antigen-specific cells of interest, but is confounded by the chronicity of infection and pathological processes. We have addressed these limitations by profiling immune responses at the site of the tuberculin skin test (TST)^17,9,18,19^. Tuberculin is a standardised clinical grade preparation of purified protein derivative (PPD) from Mtb. Inflammatory induration at the site of the TST after 2-3 days has been used extensively as a classical model of delayed type hypersensitivity dependent on T cell priming, and therefore a measure of T cell memory for Mtb antigens contained in PPD. We have previously used this model to quantify T cell recruitment and function, and to reveal exaggerated IL17 activity associated with active TB disease which could not be detected in blood^9^. The clonal repertoire of the T cell response to Mtb challenge *in vivo* has not previously been systematically evaluated. The extent to which these responses converge onto dominant T cell clones, and whether these are generalisable or idiosyncratic within a population are not known.

We addressed these questions by TCR sequencing of biopsies from the site of the TST in a cohort of 223 individuals. This approach provided us with a sensitive unbiased quantitation of T cell clones recruited and expanded in response to a standardised *in vivo* challenge. To evaluate convergence of the response to immunodominant epitopes, we developed Metaclonotypist, a modular bioinformatics pipeline for the grouping of TCR sequences based on sequence similarity. Using this pipeline, we discovered a dominant immune response to TB driven by highly public HLA-associated TCR metaclonotypes, which we expect to be a valuable resource for future biomarker discovery and reverse epitope discovery efforts in tuberculosis.

## Results

### Transcriptome-wide evaluation of maturation of the immune response to TST from day 2 to day 7

Inflammatory induration in the TST is maximal at 2-3 days, but previous flow cytometric evaluation of T cells at the site of the TST reported maximal accumulation of antigen-specific T cell responses at 7 days^20^. Therefore, we investigated the evolution of the T cell response by bulk RNAseq and TCRseq in day 2 and day 7 TSTs (Supplementary Figure 1). We recruited healthy volunteers with evidence of peripheral blood Mtb-reactive T cells, to undergo a TST in each arm (Table 1). The TST site was sampled on day 2 at one site and on day 7 at the contralateral site. Genome-wide TST-response transcriptomes at day 2 and day 7 were defined by differential gene expression compared to transcriptomes from the site of control saline injections performed in a separate set of volunteers. The TST-response transcriptomes at each time point were used to infer activity of immune response pathways at the level of cytokines, receptors, kinases and transcription factors. Both day 2 and day 7 TST transcriptomes showed activation of a comparable repertoire of canonical immune signalling pathways (Supplementary Figure 2A-B). We next identified gene expression modules associated with individual upstream regulators, which were significantly upregulated in integrated data from day 2 and day 7 TST-response transcriptomes. We found that most module expression decreased between day 2 and day 7 (Supplementary Figure 2C) consistent with homeostatic resolution of inflammatory changes. A small number of modules showed higher expression at day 7. These were all identified as being regulated by transcription factors known to be involved in cell cycle regulation (Supplementary Figure 2D).

**Table 1.**
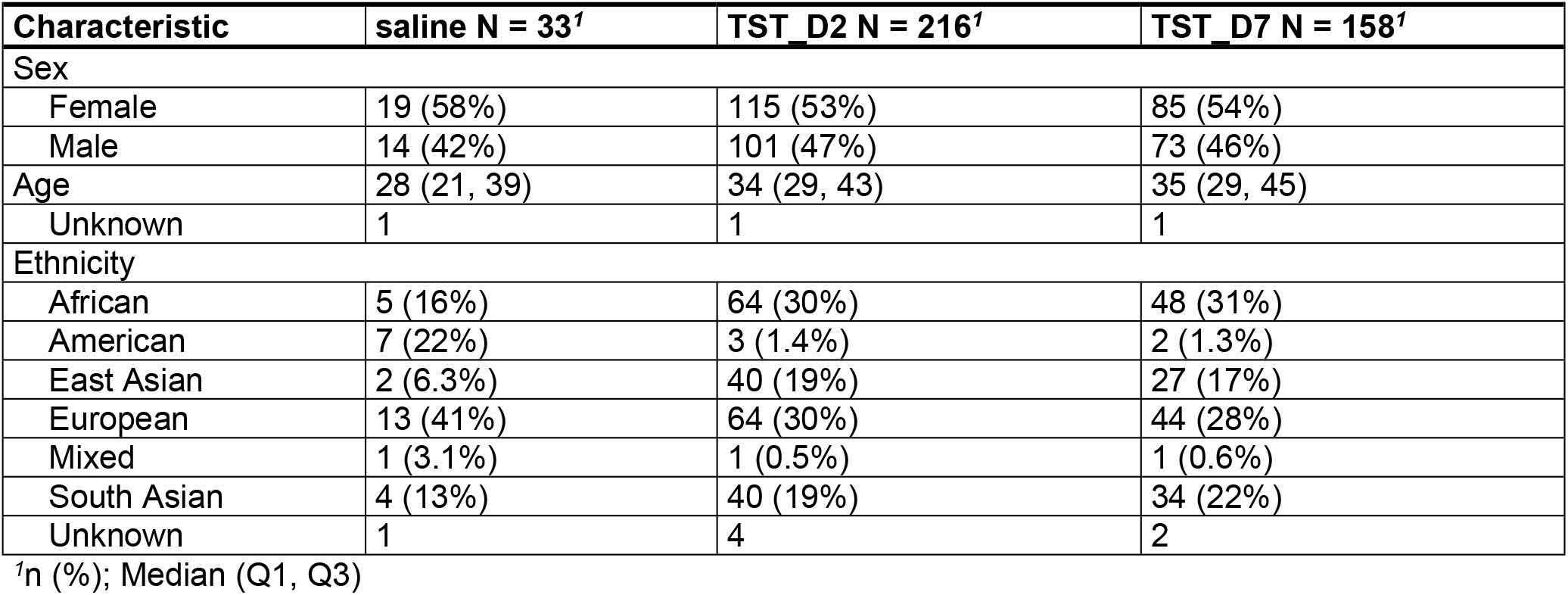
Participant overview for RNA sequencing.

Direct comparison of day 2 and day 7 TST-response transcriptomes also revealed differences in expression at the level of individual genes (Supplementary Figure 2E). Pathway enrichment analysis of the differentially expressed genes indicated increased cell cycle and mitotic activity in the day 7 TST (Supplementary Figure 2F), suggestive of increased cell proliferation at this time point. Therefore, we tested the hypothesis that the evolution of transcriptional changes between the day 2 and day 7 TSTs reflected T cell proliferation by quantifying the correlation between independently derived gene expression modules for cellular proliferation and selected T cell and non-T cell subsets. By comparison to day 2 profiles, the transcriptomes from day 7 TSTs showed significantly higher expression of the modules for cell proliferation (Figure 1A), pan-T cell, CD4 T cells and NK cells, but not modules for CD8 T cells or myeloid cells (Figure 1B). The cell proliferation module showed greatest correlation to the pan-T cell and CD4 T cell modules (Figure 1C).

**Figure 1.**
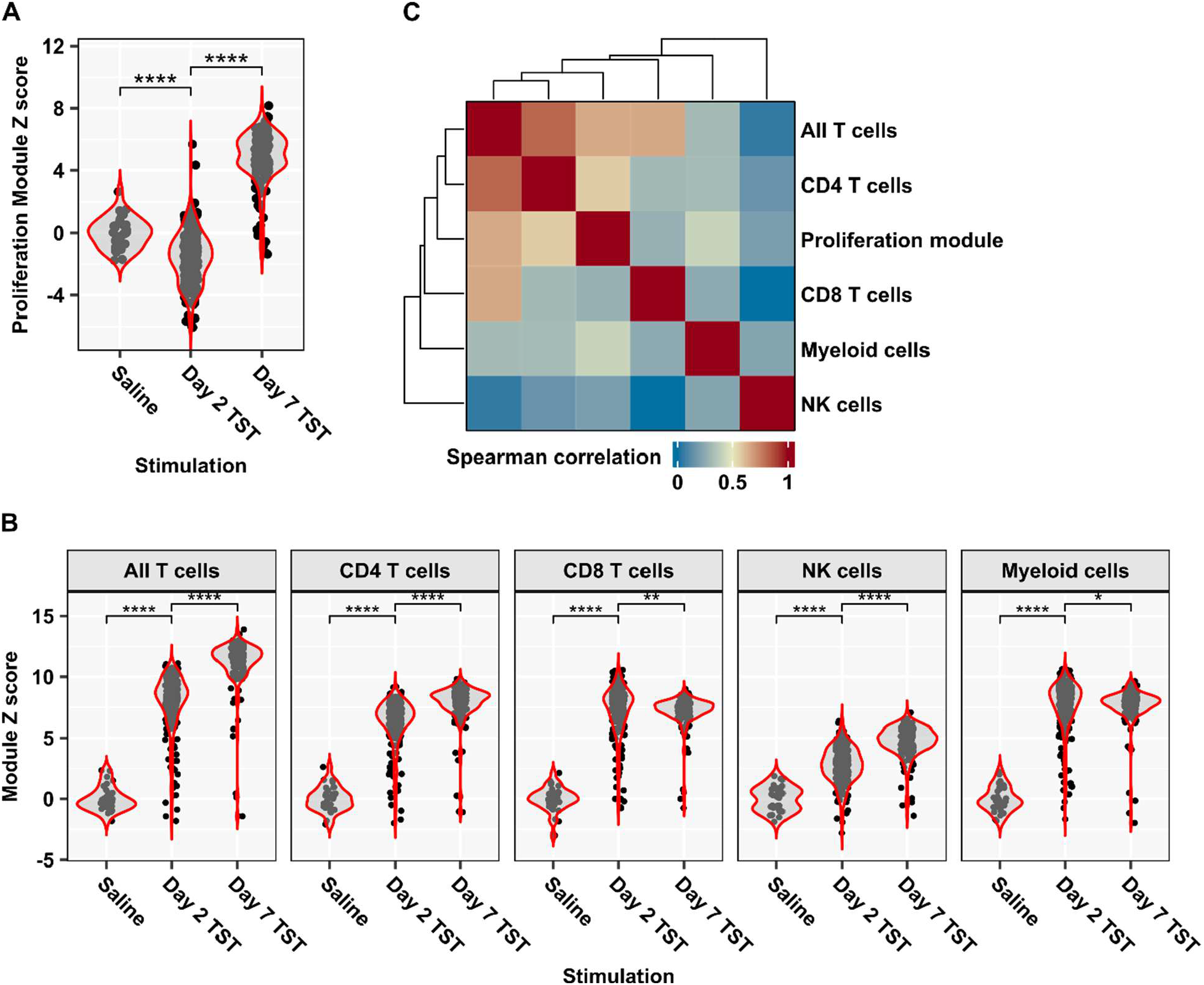
Proliferation response in Day 7 TST is correlated with CD4 T cell gene signature. Expression of cellular proliferation **(A)** and cell type-specific **(B)** modules in bulk RNA sequencing data from saline-injected control skin, Day 2 and Day 7 TSTs, shown as Z-score scaled TPM expression using saline samples as control group (n=33 saline, n=216 Day 2 TST, n=158 Day 7 TST). Wilcoxon test with multiple testing correction: * FDR<0.05, ** FDR<0.01, **** FDR<0.0001. **(C)** Heatmap of Spearman correlation matrix between gene signatures in Day 7 TSTs. Dendrogram depicts average linkage clustering of correlation coefficients.

### Evolution of reduced T cell clonal diversity in the TST

To study the nature of the T cell proliferation response revealed by our transcriptional analysis, we tracked the temporal evolution of the T cell clonal repertoire in the TST by TCRα and TCRβ sequencing of bulk RNA from day 2 and day 7 TSTs (Table 2, Supplementary Figure 1). We compared TCR repertoire diversity metrics to those of unstimulated peripheral blood samples from the same population of study participants (Figure 2A). We display metrics for the TCR β-chain, which is more diverse and informative about TCR antigen specificity^21^, but found concordant results for metrics calculated on TCR α-chain repertoires (Supplementary Figure 3A). Since repertoire diversity metrics are affected by sequencing depth (Supplementary Figure 3B-C), we down-sampled repertoires to the same size prior to this analysis. Compared to blood, the day 2 TST repertoire had an increased frequency of TCR sequences with >1 copy and a greater inequality of clone sizes as measured by Gini index, indicative of recruitment of expanded memory T cell clones. Correspondingly, the number of unique T cell clones (Richness) was reduced compared to blood. In contrast, Shannon diversity did not differ significantly from blood at day 2 and Simpson diversity even slightly increased, indicative of limited clonal dominance at this timepoint. Day 7 TSTs showed still higher proportions of expanded T cell clones, further reduced richness and Shannon diversity, and were characterised by the emergence of clonal dominance in the T cell repertoire, measured as decreased Simpson diversity and increased Gini index. Taken together, these findings suggest that the selection of T cell clones recruited to the day 2 TST is not particularly stringent, but repertoires evolve towards oligoclonality as a result of selective CD4 T cell proliferation and clonal expansion.

**Figure 2.**
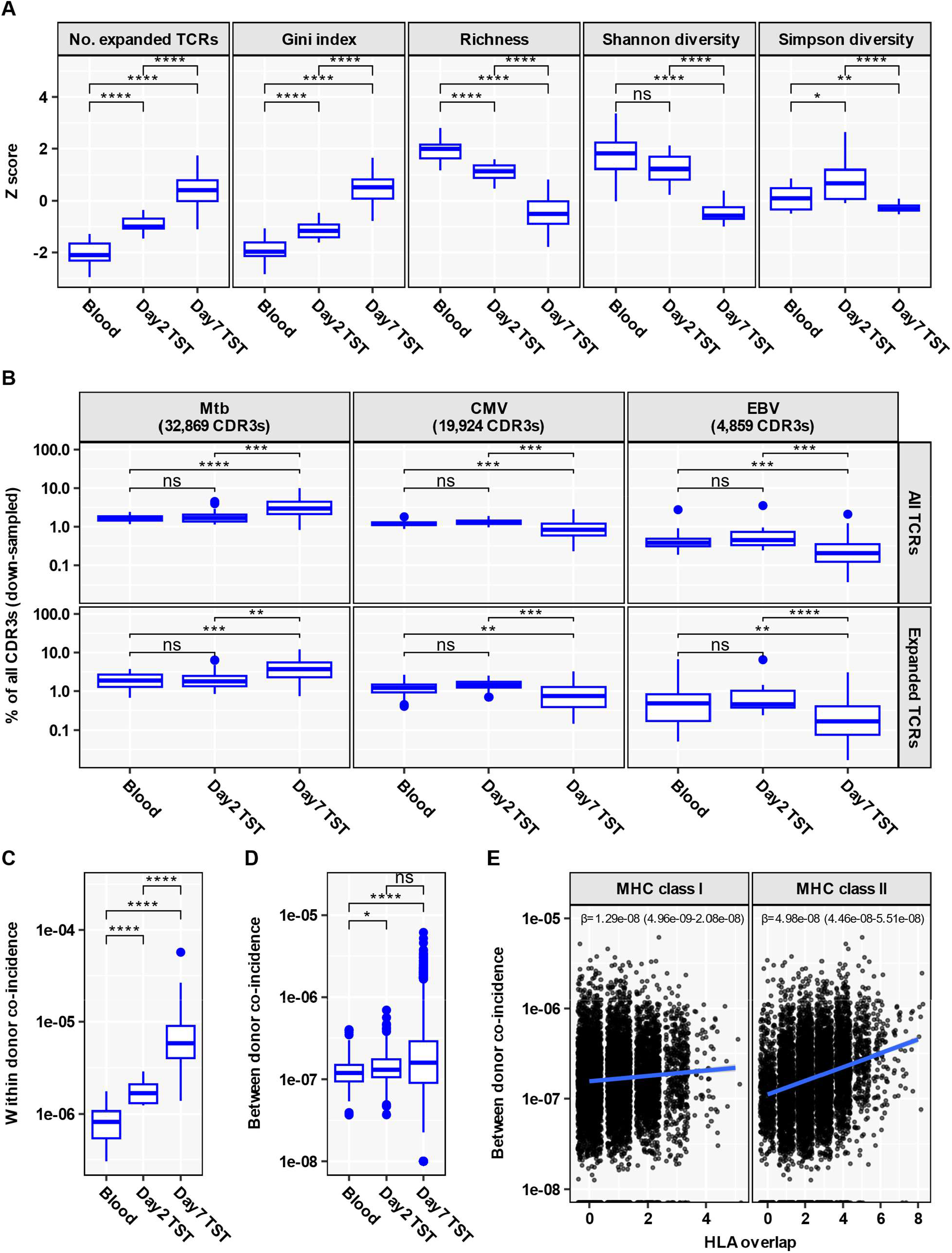
Functional restriction of the TCR repertoire in Day 7 TST yet limited inter-individual TCR sharing. **(A)** TCRβ repertoire diversity metrics, shown as Z-score values scaled across all samples. No. expanded TCRs = number of TCR sequences present more than once. **(B)** Abundance of published antigen-reactive CDR3 sequences (specific for Mtb, CMV or EBV; collated from VDJdb and McPAS databases as well as Musvosvi et al.^15^), shown as percentage of all TCRs or of all expanded TCRs (present more than once). The number of distinct published antigen-reactive CDR3s available to assess enrichment of antigen reactivity in blood and TST samples is indicated. **(C)** Within-donor convergence of distinct clones as identified by nucleotide sequence identity onto identical amino acid sequences. **(D)** Cross-donor TCR convergence, calculated between any two individuals, resulting in n=190 (Blood), n=120 (Day 2 TST) and n=8128 (Day 7 TST) pairwise comparisons. Individual β-chain bulk TCR repertoires (n=20 Blood, n=16 Day 2 TST, n=128 Day 7 TST) were down-sampled to 16,000 TCRs. Boxplots in A-D depict median and inter-quartile range (IQR), with outlier data points (more than 1.5*IQR beyond the box hinges) shown as dots. Statistical significance was assessed with Wilcoxon tests and corrected for multiple testing (ns FDR>0.05, * FDR<0.05, ** FDR<0.01, **** FDR<0.0001). **(E)** Cross-donor TCR convergence in Day 7 TSTs, stratified by the number of class 1 or class 2 HLA alleles shared between any two individuals. Each dot represents a pairwise comparison (n= 8128). The linear regression line is shown in blue, with regression coefficient (slope β) and its 95% confidence interval indicated.

**Table 2.**
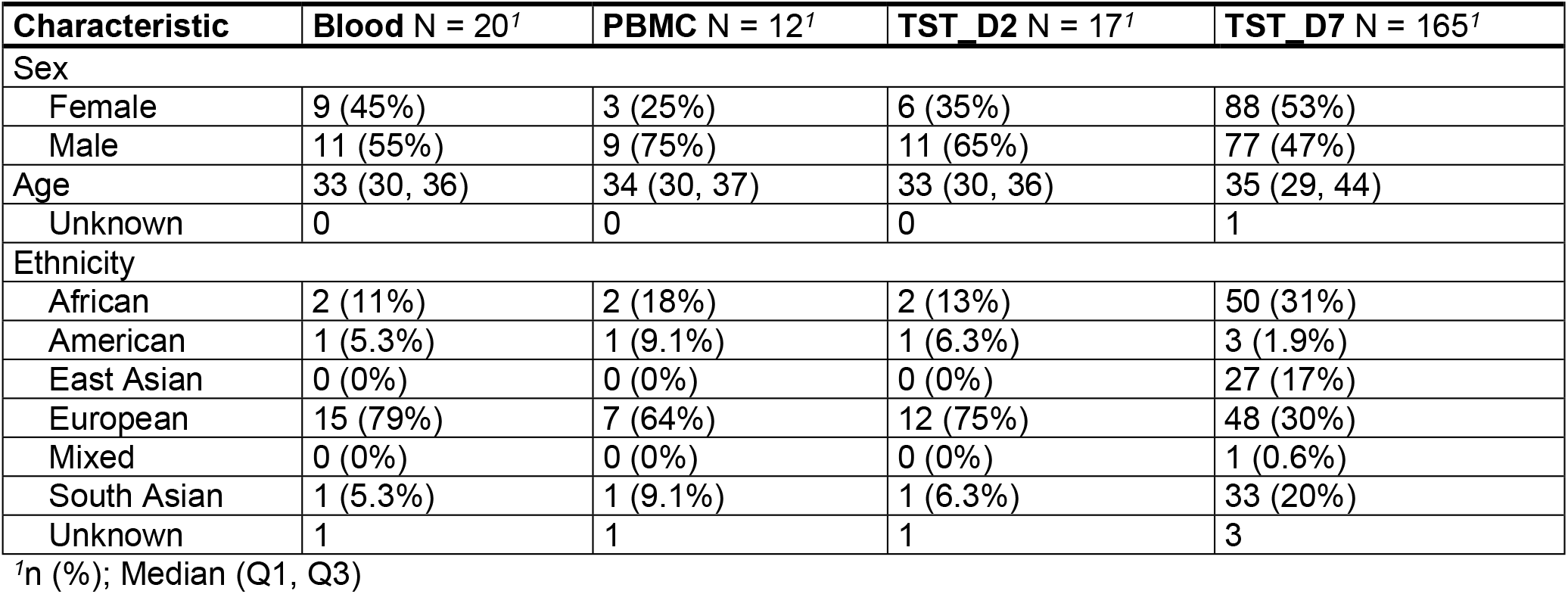
Participant overview for TCR sequencing.

### Day 7 TST is highly enriched for expanded Mtb-reactive T cell clones

Since inflammatory responses in the TST are dependent on Mtb-reactive T cells, we tested the hypothesis that the day 2 and day 7 TST TCR repertoires are selectively enriched for Mtb-reactive T cells compared to blood samples from the same individuals. We tested this hypothesis first by evaluating enrichment of previously described Mtb-reactive CDR3 sequences. We compared the TST CDR3 β-chain sequences to published CDR3 sequences derived from T cells with known antigen reactivity, including 32,869 reported Mtb-reactive TCRs identified by virtue of pMHC specificity or upregulation of T cell activation markers on ex vivo stimulation with Mtb^15,22,23^. Compared to blood, we found no enrichment for Mtb-reactive CDR3s among all TCRs or all expanded TCRs in the day 2 TST (Figure 2B, Supplementary Figure 4). However, there was statistically significant enrichment of published Mtb-reactive CDR3 sequences in day 7 TSTs compared to blood and to day 2 TSTs. As a comparison we similarly calculated the enrichment of published CMV or EBV-reactive CDR3 sequences. Day 7 TSTs showed statistically significant depletion of both CMV and EBV-reactive CDR3 sequences compared to blood and day 2 TSTs, consistent with their replacement with expanded Mtb-reactive sequences.

Next, we reasoned that antigen-driven selection of T cell responses in the TST would lead to increased functional convergence of T cell clones onto common CDR3 amino acid sequences^24^. This convergent sequence evolution was clearly evident within the repertoires of individual participants, which showed a progressive increase from blood to day 2 and then day 7 TST in coincidence probabilities (Figure 2C, Supplementary Figure 5A-C). Inter-individual analysis showed a more modest increase in average coincidence probability across day 2 TSTs from pairs of donors compared to blood, and no further significant increase in average coincidence probability among pairs of day 7 TSTs (Figure 2D, Supplementary Figure 5D-F). Interestingly, the variance of between donor coincidences increased substantially at Day 7, compatible with differential skewing of the Mtb-reactive T cell repertoires among different individuals driven by diversity in MHC-restricted antigen presentation to T cells. To further test this hypothesis, we analysed how probabilities of inter-individual coincidence in day 7 TST TCR sequences depend on HLA-allele sharing between pairs of individuals (Figure 2E, Supplementary Figure 5G-I). We found that pairs of individuals sharing multiple MHC class 2 or class 1 alleles had substantially more similar TST D7 repertoires. The dependence of repertoire overlap on HLA similarity was four-fold stronger with MHC class 2, consistent with a predominantly CD4 T cell response in the TST.

In view of the potential for inter-individual diversity of the Mtb-reactive TCR repertoire, we reasoned that evaluation of day 7 TSTs using published Mtb-reactive CDR3 sequences may substantially underestimate the enrichment of Mtb-reactive T cell clones because this analysis is inherently restricted to public TCRs. Therefore, we experimentally validated Mtb-reactive TCRs at the level of individual participants. We sequenced TCRs of peripheral blood mononuclear cells (PBMC) from a sample of the study population, following ex vivo stimulation for 6 days with PPD, or tetanus toxoid as antigen control, and selected all the CDR3s which expanded 8 fold or more in the PPD stimulated cultures but not the control unstimulated cultures^25,26^. The expanded PPD-reactive CDR3 sequences showed limited publicity among the sub-sampled study participants (Figure 3A-B, Supplementary Figure 6A-B). We then looked for overlap between the CDR3 sequences of in vitro expanded T cells and the TST repertoires from the same patient. There was no enrichment of ex vivo PPD-expanded CDR3s in the day 2 TST repertoires compared to unstimulated blood (Figure 3C-D, Supplementary Figure 6C-H). However, we found significantly greater enrichment of ex vivo PPD-expanded CDR3s in day 7 TSTs than in unstimulated blood repertoires or in day 2 TST repertoires. This pattern remained the same whether overlap was calculated for total CDR3 sequences or for unique CDR3 sequences (Figure 3C-D, Supplementary Figure 6C-H). This enrichment of Mtb-reactive CDR3s in day 7 TSTs was further increased among the most expanded TCRs. No similar enrichment of ex-vivo TT-expanded CDR3s was observed. Additionally, the odds ratio (OR) for the overlap between ex vivo PPD-reactive CDR3s was substantially greater among CDR3s which significantly expanded between day 2 and day 7 TSTs, compared to non-expanded CDR3 sequences (Supplementary Figure 7). Taken together, these results indicate that antigen non-specific accumulation of T cells in the day 2 TST is largely replaced by expanded Mtb-reactive T cell clones by day 7 post-TST.

**Figure 3.**
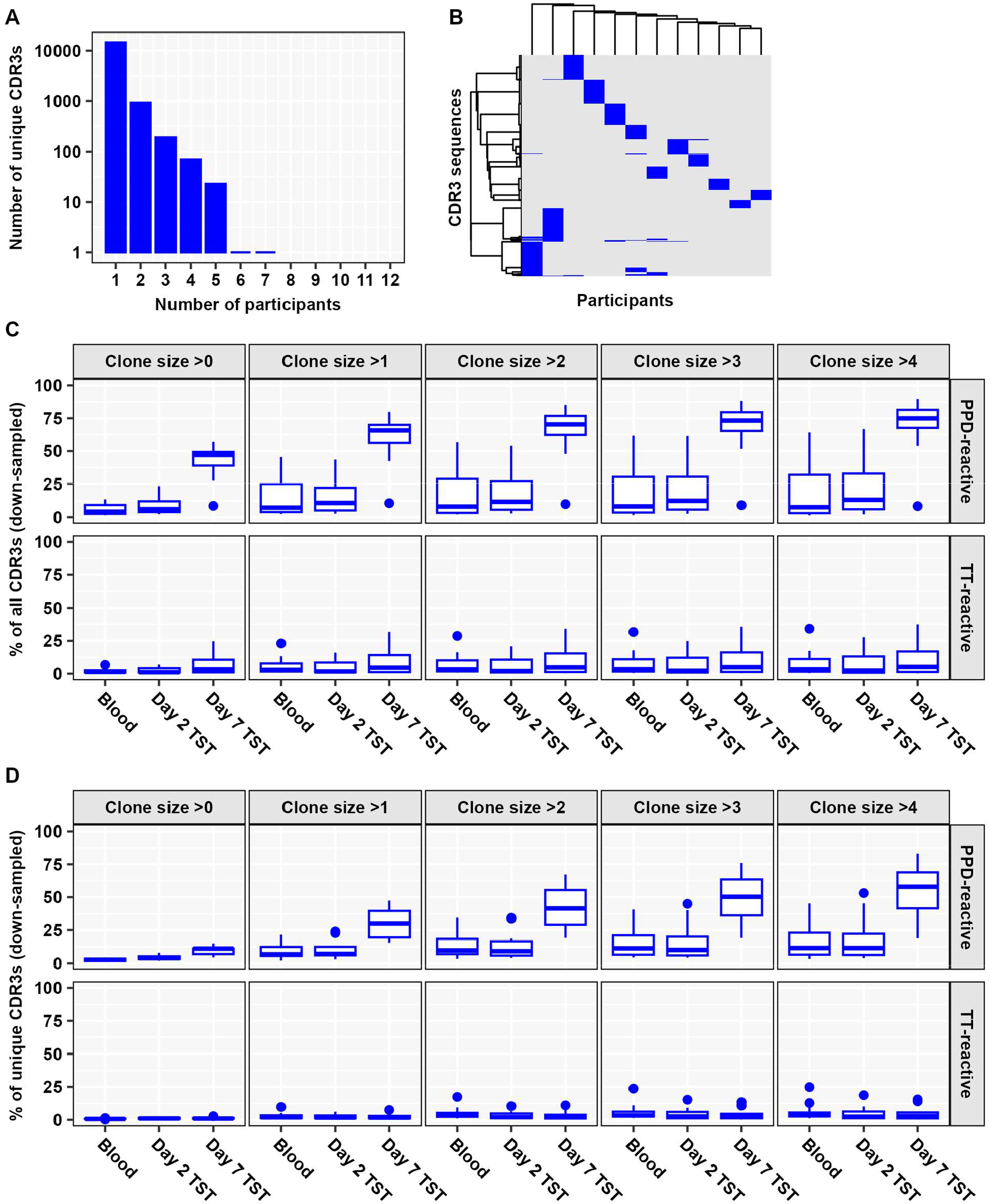
Expansion of diverse Mtb-reactive TCRs in Day 7 TST. **(A)** Histogram of number of unique PPD-reactive β-chain CDR3s shared by different numbers of participants, in ex vivo PPD-stimulated PBMC from a subset of the study population (N=12). **(B)** Heatmap of unique ex vivo PPD-reactive β-chain CDR3s (clustered by Ward D2 linkage). Abundance of total **(C)** or unique **(D)** ex vivo PPD- or TT-reactive β-chain CDR3 sequences in blood (n=12), Day 2 TST (n=11) and Day 7 TST (n=10). Individual β-chain repertoires from blood and TSTs were down-sampled to 16,000 total TCRs and stratified by clone size (= TCR count). The boxplots display median and inter-quartile range (IQR), with outlier data points (more than 1.5*IQR beyond the box hinges) shown as dots.

### Identification of Mtb-reactive metaclones in the day 7 TST

Identification of T cell metaclones, defined by similar but non-identical CDR3 sequences, which share specificity for the same peptide-MHC, can address the limitations of interindividual TCR sequence diversity, and enable antigen-agnostic identification of generalisable T cell responses to specific pMHC targets^24,27,28^. The identification of metaclones involves clustering of TCRs by sequence similarity, followed by a test for HLA-association across a cohort (Figure 4A). A number of approaches to clustering of TCR sequences have been proposed. Among these, the GLIPH2 algorithm has already been used to identify HLA-restricted Mtb-reactive T cell metaclones defined by sequence motifs^15,28^. However, selecting an appropriate clustering method for a given dataset remains a challenge, with no broad consensus on best practices^29,30^. A key difficulty lies in evaluating clustering performance, particularly in balancing the trade-off between sensitivity (for clusters often measured by retention) and positive predictive value (for clusters typically assessed as purity). Effective benchmarking of clustering algorithms, therefore, requires comparing the maximum achievable purity at fixed levels of retention. However, many existing tools generate only a single clustering solution and lack flexibility to produce clusters at multiple resolutions.

**Figure 4.**
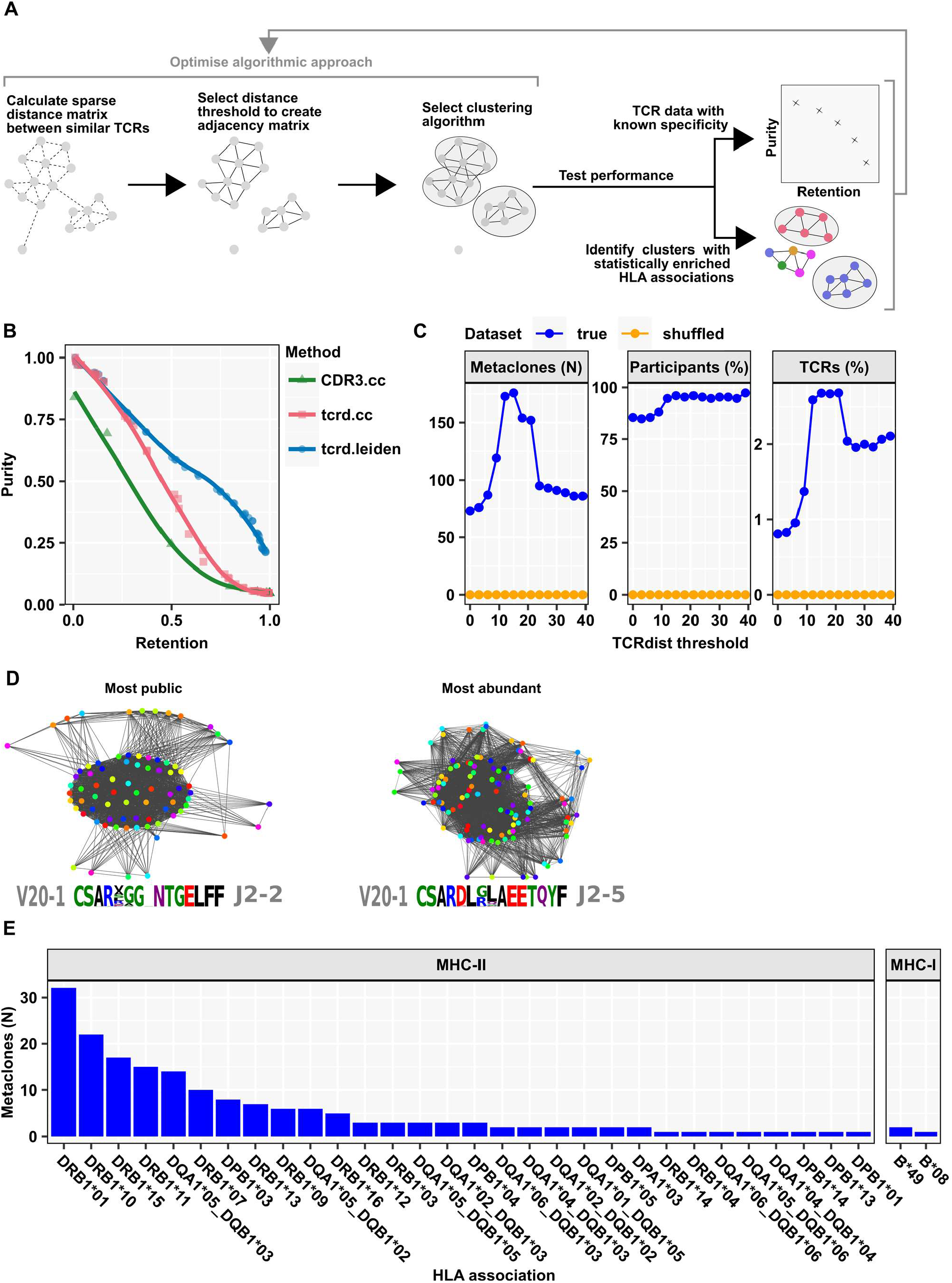
Discovery of public HLA-restricted TCR metaclones from Day 7 TSTs. **(A)** Schematic overview of the Metaclonotypist analysis pipeline and evaluation in data with known (purity/retention), or unknown specificities (by identifying significant enrichment of HLA associations, detailed in Supplementary File S4). **(B)** Trade-off between cluster purity and retention for different clustering algorithms and threshold choices benchmarked on 4840 TCRs specific to 22 distinct pMHCs from VDJdb. Metaclonotypist using TCRdist scores and Leiden clustering provides Pareto optimal clustering. **(C)** Number of HLA-enriched metaclones (left hand plot), percentage of contributing participants (middle plot) and percentage of contributing unique TCRs (right hand plot) identified at varying TCRdist thresholds by Metaclonotypist analysis of day 7 TST TCR β-chain repertoires (N=151, sub-sampled to between 5,000-10,000 TCRs per repertoire) using true or shuffled HLA allele associations. **(D)** Exemplar adjacency graphs and TCR sequence motifs of most public and most abundant day 7 TST Metaclonotypist metaclones from down-sampled repertoires (N=128 with 16,000 TCRs from each repertoire). Each node represents a single TCR stratified by distinct donors (colours). **(E)** Frequency distribution of the most significant HLA associations for each β-chain Metaclonotypist metaclones, stratified by HLA class 2 (n=177) and class 1 (n=3) allele enrichment.

To address this limitation, we developed Metaclonotypist, a modular pipeline for metaclone discovery that allows easy substitution of sequence similarity measures, thresholding choices, and clustering algorithms. Using published sets of TCR β sequences of known epitope specificity from the VDJdb database^23^, we used Metaclonotypist to systematically identify Pareto optimal algorithmic approaches (Figure 4A). Using connected component clustering, we found that adjacency graphs constructed by thresholding pairwise CDR3 edit distances were inferior to those built using TCRdist, a metric that incorporated both CDR3 and V-gene similarity (Figure 4B). At high levels of retention, we observed that Leiden clustering outperforms simple connected components clustering, as it breaks down large components into more modular, coherent clusters.

Based on these findings, we applied Leiden clustering to TCRdist adjacency graphs to analyse the day 7 TST TCR β repertoires that are highly enriched for Mtb specific T cells, but for which the specific epitopes being recognised are unknown. To ensure scalability given the large number of TCRs to be clustered in our dataset, we leveraged our previously developed symmetric-deletion lookup algorithm to rapidly identify candidate TCR neighbours in adjacency graphs^31^. We selected the TCRdist threshold which identified the largest number of TCR clusters in our dataset that were significantly enriched for individuals with a shared HLA allele (Figure 4C). The stringency of the false discovery rate control was tested by showing that random shuffling of the HLAs associated with each individual returned no HLA-enriched T cell metaclones regardless of TCRdist threshold. The identified metaclones can be represented as sequence motifs and visualised as adjacency graphs in which labelling of the individual TCR sequence by the donor reveals substantial publicity (Figure 4D). For the TCRβ repertoires, the optimal TCRdist threshold was 15, identifying 180 HLA-associated metaclones, of which 177 were restricted by HLA class II molecules. This is consistent with the known predominance of CD4 T cells in the PPD-reactive TST response, which are restricted by HLA class II. Among these, HLA-DRB1 associated metaclones were most frequent (Figure 4E).

At this threshold, >95% of study participants contributed to at least one T cell metaclone (Fig 4C, middle panel), and metaclones included TCRs from multiple different individuals (Fig 4D). However, only approximately 2.7% of unique TCR sequences were incorporated into HLA-associated metaclones, consistent with the hypothesis that the majority of Mtb specific TCRs are idiotypic (Figure 4C, right panel). We benchmarked our metaclone-discovery pipeline against clusters identified by the GLIPH2 algorithm, which passed our stringent HLA association test (Supplementary File S6-7). GLIPH2 clusters also primarily associated with class II alleles, which demonstrates the robustness of our findings with respect to the algorithmic approach. However, Metaclonotypist identified 38% more HLA-associated metaclones than GLIPH2, which accounted for 31% more unique CDR3s. This finding suggests that Metaclonotypist offers increased sensitivity to detect metaclone associations.

### Validation of Mtb-reactivity of day 7 TST derived T cell metaclones and population level immunodominance of Mtb-derived epitopes

To confirm that our day 7 TST derived, class II associated T cell metaclones represented public Mtb-reactive T cell responses, we calculated their enrichment in independent TCR sequencing data derived from people with TB compared to other diseases, or at the site of TB disease compared to blood. For internal validation, we first confirmed that the day 7 TST T cell metaclones identified by Metaclonotypist were significantly enriched in PBMC stimulated with PPD compared to PBMC stimulated with tetanus toxoid from the same study population (Figure 5A). For external validation, we then showed they were significantly enriched in peripheral blood single cell sequencing data derived from patients with TB compared to SARS-CoV-2 infection; bulk TCR sequencing of blood and lung tissue from patients with TB compared to cancer diagnoses; single cell TCR sequencing of lung tissue from patients with TB compared to cancer diagnoses; and in bulk TCR sequencing of CD4 T cells from the site of pulmonary TB disease compared to blood of the same patients (Figure 5A). We benchmarked this analysis for publicity and Mtb-reactivity of metaclones identified using Metaclonotypist against metaclones identified by the GLIPH2 algorithm. Metaclonotypist motifs showed comparable enrichment to CDR3β sequences from GLIPH2 clusters with statistically significant class II HLA-allele associations suggesting that Metaclonotypist retains similarly high specificity, despite clustering a larger proportion of TCRs into metaclones. Importantly, amino acid metaclone motifs as defined by GLIPH2 were not substantially enriched in TB samples in these datasets, indicating the need for more restrictive motif definitions. Interestingly, compared to metaclones, we found substantially lower enrichment of the full discovery set of expanded day 7 TST TCR clones, suggesting that a substantial proportion of day 7 TST TCR clones may not be specific to Mtb, and that identification of metaclones significantly improves antigen-agnostic enrichment of the Mtb-reactive T cell response.

**Figure 5.**
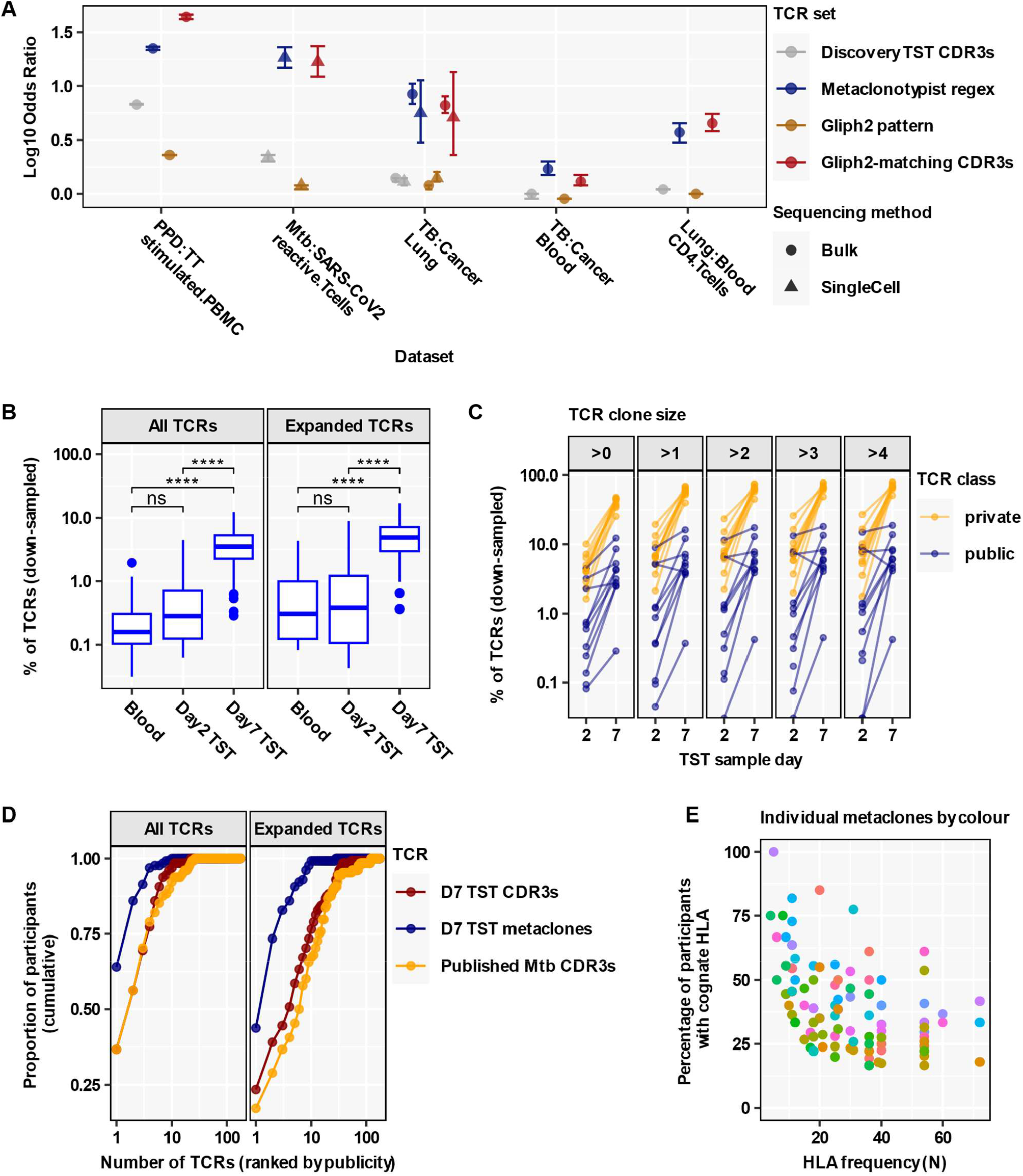
Mtb-reactive metaclones constitute a small proportion of the Day 7 TST repertoire but capture the most public response. **(A)** Relative enrichment of β-chain TCR sets derived from expanded (TCR count >1) day 7 TST repertoires (122,253 TCR clones, 151 individuals) in multiple external data sets showing point estimates and 95% confidence intervals of odds ratios (OR) in the pairwise comparisons indicated. *PBMC (bulk-TCRseq):* in vitro stimulation of PBMC from n=12 individuals with either purified protein derivative (PPD) from Mtb or tetanus toxoid (TT), followed by bulk TCR sequencing (see Figure 3), comprising n=7,743,878 PPD-stimulated and n=1,453,823 TT-stimulated TCRs. *T-cells (sc-TCRseq):* in vitro stimulation of PBMC with either Mtb lysate (n=70)^15^ or SARS-CoV2 (n=16)^49^, followed by flow cytometric sorting and single cell TCR sequencing of activated T cells, resulting in n=21,212 Mtb-reactive and n=149,208 SARS-CoV2-reactive T cells. *Lung (sc-TCRseq):* lung tissue resections from TB patients (n=5)^50^ or lung cancer patients (n=3)^51^, followed by single cell TCR sequencing, resulting in n=20,025 TB-associated and n=17,019 cancer-associated T cells. *Lung (bulk-TCRseq):* lung tissue resections from TB patients (n=13) or lung cancer patients (n=3), followed by bulk TCR sequencing, resulting in n=1,615,131 TB-associated and n=218,372 cancer-associated β TCRs. *Blood (bulk-TCRseq):* bulk TCR sequencing of whole blood samples from TB patients (n=11) or lung cancer patients (n=4), resulting in n=1,081,593 TB-associated and n=735,834 cancer-associated β TCRs. *CD4-T (bulk-TCRseq):* bulk TCR sequencing of CD4 T cells, flow-sorted from lung tissue or blood samples from TB patients (n=5), resulting in n=336,787 lung-derived and n=219,541 blood-derived β TCRs (detailed in Supplementary Table 8)^52^. **(B)** Abundance of HLA class II-restricted Metaclonotypist β-chain metaclones in TCR repertoires (each down-sampled to 16,000 TCRs; n=20 Blood, n=16 Day 2 TST, n=128 Day 7 TST), shown as percentage of all TCRs or of all expanded (>1) TCRs. Boxplots display median and inter-quartile range (IQR), with outlier data points (more than 1.5*IQR beyond the box hinges). Statistical significance was assessed with Wilcoxon tests and corrected for multiple testing (ns FDR>0.05, * FDR<0.05, ** FDR<0.01, **** FDR<0.0001). **(C)** Abundance of ‘public’ and ‘private’ Mtb-reactive CDR3s in the same individual quantified as percentage of all Day 2 or all Day 7 TST TCRs, stratified by clone size (= TCR count). β–chain TCRs were classified as ‘public’ if they matched a Metaclonotypist metaclone, or else as ‘private’ if they matched a private PPD-reactive CDR3 sequence identified from ex vivo stimulated PBMC from the same individual (see Figure 3). This analysis was restricted to individuals with paired in vitro stimulation experiments (n=11 Day 2 TST, n=10 Day 7 TST), and performed on repertoires down-sampled to 16,000 TCRs each. **(D)** HLA-class II restricted Metaclonotypist β-chain metaclones (blue), CDR3s with published Mtb reactivity (yellow), and CDR3s present in Day 7 TSTs (red) were each ranked by their publicity across 128 Day 7 TST β repertoires (each down-sampled to 16,000 TCRs) and plotted against the cumulative proportion of participants expressing the TCR. Presence of TCRs was assessed using either all TCR sequences in each sample or only expanded TCRs (present more than once). **(E)** Proportion of participants with a cognate HLA contributing to a metaclone in the discovery dataset (n=151 Day 7 TST) and frequency of the cognate HLA for each HLA class II-restricted Metaclonotypist metaclones represented by individual points.

Having confirmed that class II associated day 7 TST Metaclonotypist metaclones represented public Mtb-reactive T cells, we investigated their emergence in the TST over time. Despite some evidence for between-donor convergence of TCR sequences suggesting emergence of public T cell responses in the day 2 TST (Figure 2D), enrichment of T cell metaclones at day 2 compared to blood did not reach statistical significance (Figure 5B). In contrast, at day 7 metaclones were about 10-fold enriched in the TST samples compared to blood and day 2 TST repertoires (Figure 5B, Supplementary Figure 8). However, metaclones still only represented the minority of the total TCR sequence data at day 7. Consistent with this predominantly private response, we found no significant enrichment of previously published Mtb-reactive CDR3 sequences in the day 7 TST when these data were sub-sampled to the same number included within metaclones (Supplementary Figure 9). Therefore, despite oligoclonal restriction of the T cell repertoire over time in the TST, the day 7 repertoire may retain a substantial breadth of epitope diversity. For the subset of the study population in which we had identified Mtb-reactive CDR3s in PPD-stimulated PBMC, we compared the relative proportions of these TCRs that were private or clustered in public T cell metaclones at the level of individual participants. Private Mtb-reactive TCR sequences were more frequent than public metaclone motifs in both day 2 and day 7 TSTs across the range of TCR clone sizes (Figure 5C, Supplementary Figure 10A). There was no statistically significant difference between expansion of private and public TCRs between day 2 and day 7 TSTs, within the limitations of the sample size in this analysis (Supplementary Figure 10B-C). Despite the dominance of private TCRs at individual level, evaluation of the cumulative publicity of metaclones ranked by their publicity suggests that as few as 10 metaclones are sufficient to identify Mtb-T cell reactivity in the entire study population (Figure 5D, Supplementary Figure 11). At the level of individual metaclones there was substantial heterogeneity in publicity even among individuals with the cognate HLA allele, offering the possibility of testing Mtb-reactive T cell metaclone responses as correlates of protection and pathogenesis (Figure 5E).

## Discussion

To our knowledge, our study provides the first report of the temporal evolution of the tuberculin skin test response at the molecular level. We apply this model to establish the in vivo clonal repertoire of Mtb-reactive T cell response. At the level of individual TCRs we found this response to be almost entirely idiotypic, with very little inter-individual sharing of TCR chains even between individuals with shared HLA. Therefore, we introduce a novel analytical pipeline we have called Metaclonotypist for identification of public T cell metaclones: clusters of distinct TCRs across different individuals predicted to share peptide-MHC specificity. They enable antigen agnostic identification of T cell responses to immunodominant targets at population level by overcoming inter-individual TCR sequence diversity. We find predominantly non-antigen specific T cell recruitment to the TST initially, co-incident with peak inflammatory responses. These are then replaced by predominantly Mtb-reactive T cells through selected oligoclonal T cell proliferation. T cell metaclones derived from the day 7 TST reveal public T cell responses that are highly enriched in multiple sources of TCR sequence data from the blood and lung tissue of patients with TB compared to other diseases, and at the site of pulmonary TB compared to blood. The vast majority of TCRs enriched in the day 7 TST are not incorporated into public metaclones but remain private to individuals. Nonetheless, the cumulative publicity of the most public metaclones indicates striking population-level immunodominance of specific Mtb epitopes. This finding extends previous studies that report generalisable antigenic immunodominance at the protein level^12–14^ by providing a scalable antigen-agnostic approach to identifying generalisable immunodominance at the level of specific peptides.

The rapid accumulation of T cells at the site of the day 2 TST, which occurs at the time of maximum clinical inflammation but before evidence of cell proliferation, indicates recruitment of cells from the circulating pool. We hypothesise that local antigen causes Mtb-reactive cells to be retained at the TST site and then proliferate. Importantly, our approach therefore overcomes a limitation of peripheral blood, by sampling the TCR repertoire following recruitment from the whole in vivo T cell repertoire. Inflammatory induration in the TST is known to be dependent on prior expansion of Mtb T cell memory. Yet, we found no enrichment of public Mtb-reactive metaclones or private Mtb-reactive TCRs alongside peak inflammatory responses in day 2 TSTs compared to blood. Interestingly however, within individual donors there was a significant increase in CDR3 sequence convergence at the amino acid level in day 2 TST samples compared to blood. This finding suggests some degree of Mtb-specific T cell selection driving the TST response, albeit below the limit of sensitivity to detect through enrichment of previously reported Mtb-reactive sequences. Substantially less CDR3 convergence was evident between donors. This was partly explained by inter-individual diversity of MHC alleles but was also driven by the enormous potential diversity of the T cell receptor repertoire. Interestingly, we found a much stronger relationship of inter-individual CDR3 convergence to increased sharing of HLA class II alleles, than to increased sharing of HLA class I alleles. These data suggested that CD4 T cells are driving the maturation of T cell repertoire in the TST. This interpretation is consistent with the finding that the transcriptional signature for CD4 T cells, but not CD8 T cells, increases between day 2 and day 7 TST and correlates best with the transcriptional signature for cellular proliferation. Likewise, it is consistent with the observation that day 7 TST metaclones were almost exclusively restricted to HLA-class II alleles. Hence, we conclude that the TST is predominantly a model for CD4 T cells responses, as previously reported for T cell responses to Mtb stimulation of peripheral blood cells^15^. However, we note that this does not exclude a role for CD8 T cell responses in TB. In addition to evidence from other models that CD8 T cells may contribute to protective immunity, bulk transcriptional profiling of the TST response presented here and our recent report of single cell sequencing analysis of the TST response^32^ reveal substantial enrichment of activated CD8 T cells in the day 2 TST. The lack of further enrichment of CD8 T cell responses in the day 7 TST may be a limitation of the TST model. Nonetheless, the advantage of the skew towards CD4 T cell responses is highlighted by recent evidence that CD4 T cell responses make a more important contribution than CD8 T cells to vaccine-mediated immunity^33^.

By developing a modular computational pipeline for metaclone discovery, we were able to systematically identify algorithmic approaches with superior positive predictive value through controlled comparisons at fixed sensitivity. In combination with previously developed fast sequence similarity search tools^31^, Metaclonotypist scales accurate TCRdist-based clustering to large datasets and allows flexible optimisation of parameters. The optimised Metaclonotypist-pipeline identified significantly more HLA-associated metaclones in our dataset than the principal metaclone discovery algorithm (GLIPH2) that has been previously used to identify Mtb-reactive metaclones^15,28^, while maintaining comparable enrichment in a wide-ranging external validation. Despite this increased sensitivity, the majority of day 7 TST TCRs remained excluded from Metaclonotypist metaclones, which we have labelled as private T cell responses. We anticipate that further artificial-intelligence driven improvements in measures of TCR sequence similarity^34^ or corrections for recombination biases^27^ will allow clustering of more distant TCRs with shared peptide-MHC specificity. Therefore, our current approach provides only a minimum estimate of publicity in the Mtb-reactive T cell repertoire. Even so, as few as 10 metaclones were sufficient to identify generalisable Mtb reactive T cell responses in our study population of ≥128 individuals. Interestingly, for most metaclones not all individuals with the restricting HLA allele had detectable metaclone TCRs. This finding is compatible with potential inter-individual heterogeneity of Mtb responses to particular pMHCs, which in conjunction with future improvements in metaclone discovery and deeper TCR sequencing may allow linking T cell metaclone responses to differential outcomes of Mtb infection.

Consistent with most data on HLA class II restriction of CD4 T cell responses, we found day 7 TST metaclones to be predominantly restricted by HLA-DR alleles. This skew is attributed to the higher prevalence of diverse DR alleles, a structure that allows them to bind a wider variety of peptides, and potentially higher levels of expression than HLA-DP and HLA-DQ alleles on antigen presenting cells^35–37^. The application of bulk TCR sequencing maximised the depth of data and therefore the sensitivity of our analysis. Our primary approach focussed on β-chain sequences to leverage their greater diversity and provide greater discrimination of the TCR repertoire than possible by analysis of α chains. Reassuringly, analyses of α-chain sequences replicated the findings from β chain repertoires. In future work, identification of public metaclone motifs in single cell sequencing datasets can identify associated αβ-chain pairs. Although the more limited depth of single cell sequencing limits sensitivity for metaclone discovery, αβ-chain pairs are necessary to pursue reverse epitope discovery to test the prediction that metaclone clustered TCRs share epitope specificity, develop functional T cell assays to evaluate the association of immunodominant T cell metaclones with clinical outcomes of infection, and enable innovation in vaccine design based on specific protective epitopes in place of protein antigens.

Our application of Metaclonotypist, combining fast clustering and HLA associations, to TCR sequence data from the day 7 TST that is highly enriched for Mtb-reactive T cells recruited from the in vivo repertoire, has identified a catalogue of metaclones, which together span the majority of the study population. We hypothesise that these metaclones will provide powerful approaches to improve disease-risk stratification in Mtb-infected people, and, in combination with single-cell sequencing will enable identification of epitopes to resolve protective and pathogenic T cell immunity critical to the development of more effective vaccines.

## Methods

### Study approvals

Research ethics and regulatory approvals for the present study were provided by UK research ethics committees (reference numbers: 11/LO/1863, 14/LO/0505, and 18/LO/0680). All study participants provided written informed consent.

### Study population and sampling

Study participants comprised healthy HIV seronegative adults, 18-60 years of age, with immune memory for Mtb-specific antigens identified by positive peripheral blood IFNγ release assays using the QuantiFERON Gold Plus Test, but no clinical or radiological evidence of active tuberculosis. Peripheral blood mononuclear samples purified by Ficoll-Paque Plus (GE Healthcare Biosciences) density gradient centrifugation of whole blood, were collected on participant enrolment, and cryopreserved in foetal calf serum (FCS, Sigma) supplemented with 10% DMSO (Sigma). Blood RNA samples were collected into Tempus RNA preservative tubes (Thermo Fisher Scientific), used to extract total RNA with the Tempus Spin RNA Isolation Kit (Ambion; Life Technologies) and remove genomic DNA using the TURBO DNAfree kit (Ambion, Life Technologies). Participants then received 2U intradermal tuberculin (Serum Statens Institute) or control saline injections as previously described^9,17,19^. 3mm punch biopsies from the injection sites were collected and cryopreserved in RNALater (Qiagen) at designated time points and used to extract total RNA as previously described^9^.

### RNA sequencing and analysis

Total RNA from TSTs were subjected to genome wide mRNA sequencing as previously described^9^. This provided a median of 22 million (range 10-50 million) 41 bp paired-end reads per sample. RNAseq data were mapped to the reference transcriptome (Ensembl Human GRCh38 release 111) using Kallisto^38^. Transcript-level counts were summed on gene level, and annotated with Ensembl gene ID, gene symbol and gene biotype using the R/Bioconductor packages tximport and BioMart. Raw counts of 23,820 Ensembl gene IDs, retained after exclusion of pseudogenes, were used for differential expression analysis with the SARtools implementation of DeSeq2^39^, with a false discovery rate (FDR) <0.05 and log2 fold difference of ≥1. For all other analyses, raw counts were converted into transcripts per millions (TPM) values, and log2 transformed after the addition of a pseudocount of 0.001. Duplicated gene symbols were filtered by retaining the gene with highest expression per sample.

Upstream regulator analysis of the differentially expressed genes was performed using Ingenuity Pathway Analysis (Qiagen). This was visualised as a network diagram using the Force Atlas 2 algorithm in Gephi v0.9.4, and used to derive co-regulated gene-expression networks as previously described^40^. Briefly, this analysis was restricted to upstream regulators predicted to be significantly activated (Z-score>2, adjusted p-value<0.05), targeting at least 4 downstream genes, and annotated with one of the following functions: cytokine, kinase, transmembrane receptor, and transcriptional regulator, representing the canonical components of pathways which execute transcriptional reprogramming in immune responses. For each upstream regulator, pairwise Spearman correlations of the TPM expression values of the target genes were calculated among TST samples.

Upstream regulators were selected as ‘significant’ if the average co-correlation was significantly (FDR <0.05) greater than the distribution of average correlation coefficients obtained from 100 iterations of selecting an equivalent number of random genes. Reactome pathway enrichment of differentially expressed genes was analysed with the XGR R package^41^. For visualisation, 20 pathway groups were identified by hierarchical clustering of Jaccard indices to quantify similarity between the gene compositions of each pathway. For each group, the pathway with the largest total number of genes was then selected to provide a representative annotation.

Transcriptional modules for T cell proliferation^40^ and cell types present in the TST^32^ have been derived and published previously. Their gene composition is listed in Supplementary File S1. The expression of each module was quantified as the arithmetic mean log2 TPM value of its constituent genes.

### Ex vivo PBMC stimulation

Frozen PBMC were thawed at 37°C, washed in RPMI-1640 media (Thermo Fisher Scientific) with 10% FCS, at resuspended at 10^6^ cells/ml in RPMI with 5% heat-inactivated male human AB serum (Sigma). 2×10^5^ were seeded into individual wells of round bottom 96-well plates (Thermo Fisher Scientific) with one of 10 μg/mL purified protein derivative of Mtb (PPD; Serum Statens Institute), 100 μg/mL tetanus toxoid (NIBSC), or control buffer for 6 days at 37°C and 5% CO_2_. At the end of this incubation period, plates were centrifuged (400g for 5 minutes) and the resulting cell pellets were lysed in RLT buffer (Qiagen). Samples from triplicate wells were pooled for RNA extraction using the RNneasy kit (Qiagen). Up to 5 separate pooled samples were collected for each individual, for each stimulus.

### T cell receptor (TCR) sequencing and analysis

RNA extracted from skin samples, ex vivo stimulated PBMC and peripheral blood Tempus tubes were subjected to sequencing of TCR α and β-genes using an established quantitative TCR sequencing pipeline that integrates experimental library preparation and computational analysis with Decombinator V4^26,42,43^ which defines and quantifies a ‘TCR clone’ by its nucleotide sequence and reports its V, J and CDR3 annotation. To account for different sequencing depth between samples, repertoire metrics were calculated after downsampling all samples to 16,000 unique molecular identified (UMI) reads (Supplementary Figure 1).

TCR α and β CDR3s from whole blood or skin biopsies were annotated as CMV, EBV or Mtb-reactive, if they were listed as sequences known to target these pathogens in the VDJdb TCR repository (https://vdjdb.cdr3.net/; accessed 01/10/2024)^23,44^, the McPAS database (http://friedmanlab.weizmann.ac.il/McPAS-TCR/; accessed 16/09/2023)^22^ or in Table S2 from Musvosvi et al^15^. The collated antigen-reactive CDR3 sequences are summarised in Supplementary File S2. To identify antigen-reactive CDR3s from in vitro cultures, we identified CDR3 sequences with ≥8-fold increased abundance in antigen stimulated, but not unstimulated PBMC compared to whole blood from the same individual, as previously described^26^. CDR3s absent in matching blood were set to the median blood CDR3 abundance of 1 to allow expansion calculations for all in vitro CDR3s. To complement this analysis, we also used a more stringent definition of expanded T cell clones as previously described^40,45^. In this approach, CDR3s in Day 7 TST samples were defined as ‘significantly expanded’ if their observed abundance was greater than expected using a Poisson distribution derived from Day 2 TST counts with FDR<0.1%.

TCR repertoire diversity was assessed by the number of expanded TCR sequences (count>1), Gini index (repertoire inequality), and Hill Diversity indices. These diversity indices are defined as 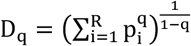 where R is the number of distinct TCRs, p_i_ the clonal frequency of the i-th clone, and q a parameter that determines the relative weight put on clonal abundance. We compared richness (total number of distinct TCRs), *D*_0_= *R*, Shannon Diversity (exponential of Shannon entropy), 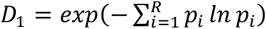, and Simpson diversity (inverse of Simpson’s index), 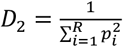. Among these measures Simpson diversity is most sensitive to clonal dominance, while Richness completely disregards variability in clonal expansions.

Within- and cross-donor convergence of TCR sequences was calculated as previously described^24^. Within-donor convergence was calculated as the proportion of all pairs of distinct clonotypes (as defined by nucleotide sequence identity) which were functionally convergent, i.e. that encode the same protein. We define *n*_i_ as the number of distinct clonotypes encoding the *i* − *th* TCR with i = 1, …, S, where *S* is the number of unique clonotypes. We also define N = Σ n_i_ as the total number of clonotypes in the sample. We then can estimate the probability of coincidence within a sample as: 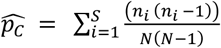. In comparing across samples, we define clonotypes by nucleotide sequence and donor identity. Defining n_l,1_ and n_l,2_ as the sampled counts of the *i* − *th* TCR in donor 1 and donor 2, respectively, we estimate the probability of cross-donor convergence using: 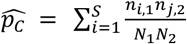, where 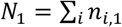 *and* 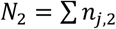 is the total number of clonotypes in the two samples.

### HLA genotype imputations

DNA from participants was extracted from cryopreserved whole blood using the QIAamp spin column (Qiagen). Genotyping was conducted using the Illumina Infinium Global Diversity Array. HLA imputation was performed on the Michigan Imputation Server^46^ using genotyped autosomal variants across the study population, filtered to include only SNPs with a minor allele frequency (MAF) of ≥5% and a call rate of ≥95%. Briefly, typed SNPs within the MHC region (6:27970031-33965553; hg19) were phased with Eagle (v2.4) and imputed using Minimac4 with the four-digit multi-ancestry HLA imputation reference panel (v2). Imputed SNPs with an imputation score (R^2^) <0.8 were excluded, resulting in high-confidence HLA alleles for 158 individuals (Supplementary File S3).

### Benchmarking TCR clustering approaches using Metaclonotypist

To compare TCR clustering approaches, we implemented Metaclonotypist, a modular computational pipeline for metaclonotype discovery, and benchmarked clustering performance using TCRs with known pMHC specificity from the VDJdb database^23^.

Metaclonotypist proceeds in a series of steps (Figure 4A). Metaclonotypist first calculates pairwise distances between TCRs according to sequence similarity metrics, from simple Levenshtein edit distances applied to the CDR3 sequence to more advanced metrics such as TCRdist. This first step can be optionally sped up by pre-filtering of candidate sequence neighbour pairs using the symmetric deletion lookup algorithm. It next generates an adjacency graph between sequences, by thresholding the pairwise sequence similarity with respect to a tuneable threshold. Each node in this graph represents a TCR found in an individual’s repertoire, and edges connect all nodes with a similarity below the threshold. Within the graph Metaclonotypist then identifies putative metaclones by clustering. Clustering is performed using community detection algorithms as implemented in igraph^47^. Importantly, each step is modular and support multiple choices to allow benchmarking of alternative approaches using different sequence similarity metrics, threshold choices for adjacency graph construction, and clustering algorithms.

To construct a benchmarking task, we selected data from all pMHCs with at least 220 associated TCR β sequences from VDJdb following filtering and data standardisation using tidytcells. We then randomly down-sampled TCR repertoires from pMHCs with a greater number of sequences to obtain a dataset of 4840 TCR β sequences equally balanced across 22 pMHCs.

Clustering involves a multi-objective optimisation, with ideal clustering having both high purity and retention. To allow controlled comparisons across similarity metrics and clustering algorithms, we systematically varied distance thresholds for each method to be able to identify Pareto optimal solutions. We defined cluster purity as the weighted average of the dominant class frequency in each cluster: 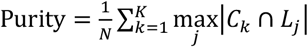, where N is the total number of TCRs, K the total number of clusters, C_k_ the set of TCRs associated with cluster k, and L_j_ the set of TCRs associated with label j (here representing a specific epitope). We defined clustering retention as the fraction of all TCRs assigned to non-singleton clusters: 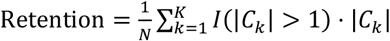, where *I*(|*C*_k_| > 1) is an indicator function that is one if |*C*_k_| > 1 and 0 otherwise.

Using this benchmarking approach, we compared connected component clustering of adjacency graphs based on CDR3 Levenshtein distance, which simply groups all TCRs connected by at least one edge into a cluster, to more advanced algorithms. Our results suggest that the more advanced TCR sequence similarity metric TCRdist is superior to simple Levenshtein distance calculated on the CDR3 alone. We furthermore found that Leiden clustering, which breaks up large connected components into multiple clusters where this increases cluster modularity, maintains higher purity at larger thresholds.

### Discovery of HLA-associated metaclones with Metaclonotypist

We considered metaclones as a set of β-chain TCR clones with an imputed common HLA-peptide specificity^27^. We identify putative metaclones by clustering TCRs based on sequence similarity and testing the HLA association of TCR clusters. We combined bulk-sequenced Day 7 TST repertoires for β chains. To reduce uneven sampling, we down-sampled large repertoires to 10,000 total counts and excluded repertoires with <5,000 total counts. To increase confidence of restricting metaclone discovery to Mtb-reactive TCRs, only those with count >1 in these down-sampled repertoires were selected for analysis. TCRs with CDR3 amino acid length ≤5 were excluded from analysis. We then identified all pairs of TCRs that differ by ≤2 edits in their CDR3 hypervariable region using the symmetric deletion lookup algorithm^31^. We next calculated TCRdist scores between these pre-pruned TCR pairs using TCRdist3^27^. Based on our preliminary benchmarking we used Leiden clustering for our identification of metaclonotypes in the day 7 TST (with parameters: resolution=0.1, objective_function=‘CPM’, n_iterations=4). After examining the effect of varying thresholds, we represented the TCR repertoire as an undirected graph based on the sparse adjacency matrix obtained by thresholding TCRdist scores ≤15.

Each cluster was tested for HLA association, by comparing the expression of specific HLA alleles by Fisher’s exact test between two groups of individuals: those contributing at least one TCR to a cluster and the remainder of the population. Associations for HLA class II alleles (DP, DQ, DR) and class I alleles (A, B, C) were tested separately. HLA-association of metaclones with the DQ locus was tested with respect to all potential DQ heterodimers, by combining DQ alleles for the α and β HLA chain to account for the highly polymorphic nature of both the α and β chain of HLA DQ. P values were corrected for multiple testing using the Bonferroni-Hochberg procedure at a False Discovery Rate (FDR) of 0.1 where the number of tests was set equal to the product of the number of tested clusters and times the number of tested HLA alleles. To limit multiple testing, we only assessed association of clusters containing TCRs from ≥4 individuals with HLAs found in ≥4 individuals across the population. As a control, the link between HLA haplotype and individuals was randomly shuffled.

### Metaclone visualisation

Sequence logos were constructed in python, using the seqlogos_vj plotting submodule of the pyrepseq package^48^. Graphs of TCR sequence similarity within a metaclone were visualised using Python bindings to the igraph package. Each node represents a TCR, and distinct colours are used to indicate donor origin. Nodes are connected by unweighted edges whenever corresponding TCRs were below the threshold of sequence similarity used for metaclone discovery.

### GLIPH2 analysis

GLIPH2 analysis was undertaken on the same set of Day 7 TST β-chain repertoires as described for metaclonotype discovery above, using default settings and CD48_v2.0 reference. The Metaclonotypist approach was then mirrored, by selecting only GLIPH2 similarity clusters containing TCRs from ≥4 individuals and testing for associations with HLAs found in ≥4 individuals. Each cluster was tested for HLA association by Fisher’s exact test with FDR <0.1, testing separately associations with class I and class II HLA alleles.

### Quantification of Metaclonotypist and GLIPH2 metaclones

Supplementary Files S4 and S5 list the HLA class II and class I restricted β metaclones respectively, as identified by Metaclonotypist from day 7 TST samples. The tables include the restricting HLA allele for each metaclone (defined as the most significant HLA association), as well as the associated V gene(s) and a regular expression for the clustered CDR3 amino acid sequences. To identify and quantify β-chain TCR sequences from various datasets that match a pre-defined class II associated metaclone in the context of the correct TCR chain, the V gene and CDR3 of each TCR was compared against the V gene and CDR3 regular expression of each metaclone.

Supplementary Files S6 and S7 list the HLA class II and class I restricted β chain GLIPH2 clusters, respectively, identified from day 7 TST samples. The tables include the restricting HLA allele (defined as the most significant HLA association), as well as the clustered CDR3 sequences and a regular expression for the shared CDR3 pattern. β-chain TCR sequences from various datasets were identified as matching a pre-defined class II associated GLIPH2 cluster in two different ways: a) if their CDR3 amino sequence contained the regular expression of the GLIPH2 motif (GLIPH2 pattern G.T was excluded from analysis to increase specificity), b) if their CDR3 amino acid sequence was part of the GLIPH2-clustered set of CDR3 sequences.

### Datasets for external validation of TST-derived metaclones

Processed single cell TCR sequencing data from activated T cells following in vitro stimulation of PBMC from n=70 individuals with Mtb lysate were accessed from Supplementary Table S2 in the publication by Musvosvi et al^15^. Only good quality cells (flag = GOOD) were included, resulting in 21,212 cells with β-chain data.

Processed single cell TCR sequencing data from activated T cells following in vitro stimulation of PBMC from n=16 individuals with SARS-CoV2 were provided by Lindeboom et al^49^. All longitudinal samples per patient were included, resulting in 149,208 cells with β-chain data.

Single cell TCR sequencing FASTQ data from human lung of n=5 TB patients^50^ were downloaded from the NCBI gene expression omnibus resource (GSE253828) and processed with 10x Genomics CellRanger (v7.1.0) using the vdj pipeline and VDJ-T reference version 7.1. Single cell TCR data from ‘filtered_contig_annotations.csv’ output files were integrated across all patients, resulting in 20,025 cells with β-chain data.

Single cell TCR sequencing FASTQ data from human lung of n=3 lung cancer patients^51^ were downloaded from the NCBI gene expression omnibus resource (GSE154826) and processed with 10x Genomics CellRanger (v7.1.0) using the vdj pipeline and VDJ-T reference version 7.1. Single cell TCR data from ‘filtered_contig_annotations.csv’ output files were integrated across all samples (including tumour and normal lung tissue) from all patients, resulting in 17,019 cells with β-chain data.

Processed bulk TCR sequencing data for lung tissue and whole blood from TB patients and cancer controls, as well as for sorted CD4 T cells from TB lung and TB blood were provided on Adaptive Biotechnologies’ ImmunoSEQ website (https://clients.adaptivebiotech.com/). The cohort has been previously described^52^, and an overview of utilised files and their metadata is provided in Supplementary File S8. Only functional TCR sequences were included (sequenceStatus = In), and the vMaxResolved column was used as V gene annotation, but with the allele information excluded (e.g. ‘TCRBV06-01*01’ became ‘TCRBV06-01’). Since the ImmunoSeq naming of TCR V genes differs from the IMGT nomenclature used for metaclone definitions, V gene names were made compatible prior to searching for metaclone matches. This included, within the ImmunoSeq annotations, replacement of ‘TCRBV’ with ‘TRBV’, and the removal of leading zeroes from V gene alleles (e.g. replacing TRBV06-06 with TRBV6-6). β-chain data were available for the lung dataset (n=13 TB patients and n=3 cancer controls), the blood dataset (n=11 TB patients and n=4 cancer controls), and the CD4 T cell dataset (n=5 TB patients). All samples per patient were included, and data integrated across disease and tissue groups, resulting in n=1,615,131 TB-associated and n=218,372 cancer-associated β TCRs for the lung dataset; n=1,081,593 TB-associated and n=735,834 cancer-associated β TCRs for the blood dataset; and n=336,787 lung-derived and n=219,541 blood-derived β TCRs for the CD4 T cell dataset.

### Statistics and data visualisation

Analyses were performed in R (version 4.3.3) or Python (version 3.10.4). Data were visualised and figures assembled using R’s tidyverse and ggpubr packages. Statistical differences were assessed using the tests and significance thresholds stated in the text and figure legends. Wilcoxon tests with FDR correction for multiple testing were performed with the wilcox_test or pairwise_wilcox_test functions from the rstatix package in R. Base R functions cor() and lm() were used for Spearman correlation and linear regression analyses, respectively, with confint() to calculate confidence intervals for regression coefficients. R packages pheatmap and ComplexHeatmap were used to create heatmaps. Odds ratios and their confidence intervals were calculated with the fisher.test() function from the stats R package. Metaclones were identified using Metaclonotypist (written in Python) and visualised as described above.

## Data and code availability

At the time of peer-reviewed publication, all analysis code will be made available on GitHub; raw RNA sequencing data in FASTQ format will be available under controlled access at the European Genome-Phenome Archive with accession number EGAD50000001208 (https://ega-archive.org/); processed RNAseq data will be available at Array Express with accession number E-MTAB-14687 (https://www.ebi.ac.uk/arrayexpress); raw TCR sequencing data in FASTQ format will be available at NCBI Short Read Archive with accession number PRJNA1208718 (https://www.ncbi.nlm.nih.gov/sra); processed TCR sequencing data will be available from UCL’s Research Data Repository (https://rdr.ucl.ac.uk/; DOI 10.5522/04/28049606)

## Footnotes

## Acknowledgements

This work was supported by Wellcome Trust awards to MN (207511/Z/17/Z and 306550/Z/23/Z). MN and GP acknowledge support from NIHR Biomedical Research Funding to University College London Hospitals. ATM acknowledges support by the Royal Free Charity. JCK acknowledges support from NIHR Oxford Biomedical Research Centre. JJ is funded by the Deutsche Forschungsgemeinschaft (DFG, German Research Foundation, 531855214). We thank Michelle Berin, Zandile Maseko and Kimberlee Gunn, for supporting participant recruitment and sampling.

## Author contributions

Conceived and designed the study: CT, ATM, BMC, MN

Sample and clinical data collection: MB, RBM, SC, ML, HK, SL, PO, GP, AL, MN

Laboratory analysis: CT, JR, AC, IU, GN, SB, RBM, GP

Data analysis: CT, ATM, RS, PZ, JJ, AK, FP, WS, VR, GP, JCK, BMC, MN

Manuscript preparation: CT, ATM, BMC, MN with input from all authors

