## Supplementary Figures for "Evolution of T cell responses in the tuberculin skin test reveals generalisable Mtb-reactive T cell metaclones"

### Supplementary data contents

**Supplementary Figure 1**

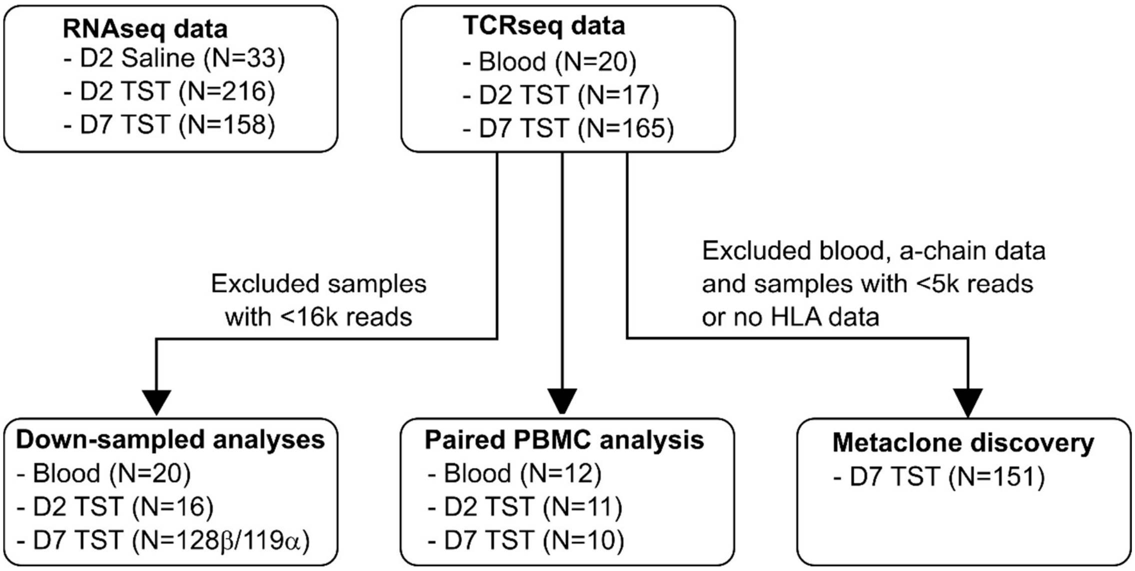

***Consort diagram for application of samples in RNAseq and TCRseq analyses.***

The data analysed in the present manuscript was derived from a cohort 216 individuals who underwent an intradermal tuberculin skin test in each arm and a separate cohort of 33 individuals who underwent an intradermal saline injection in one arm. In the TST cohort, total RNA from punch biopsies at the site of the TST were available from all 216 individuals at day 2, and 158 individuals at day 7. In the 'saline' cohort, total RNA from punch biopsies at the site of saline injection was available from all 33 individuals at day 2. Genome-wide transcriptional profiling by RNA sequencing was performed in all available RNA samples from these cohorts. TCR sequencing initially focussed on a subset of day 2 TST and paired blood samples. This was then extended to all day 7 TST samples.

**Supplementary Figure 2**

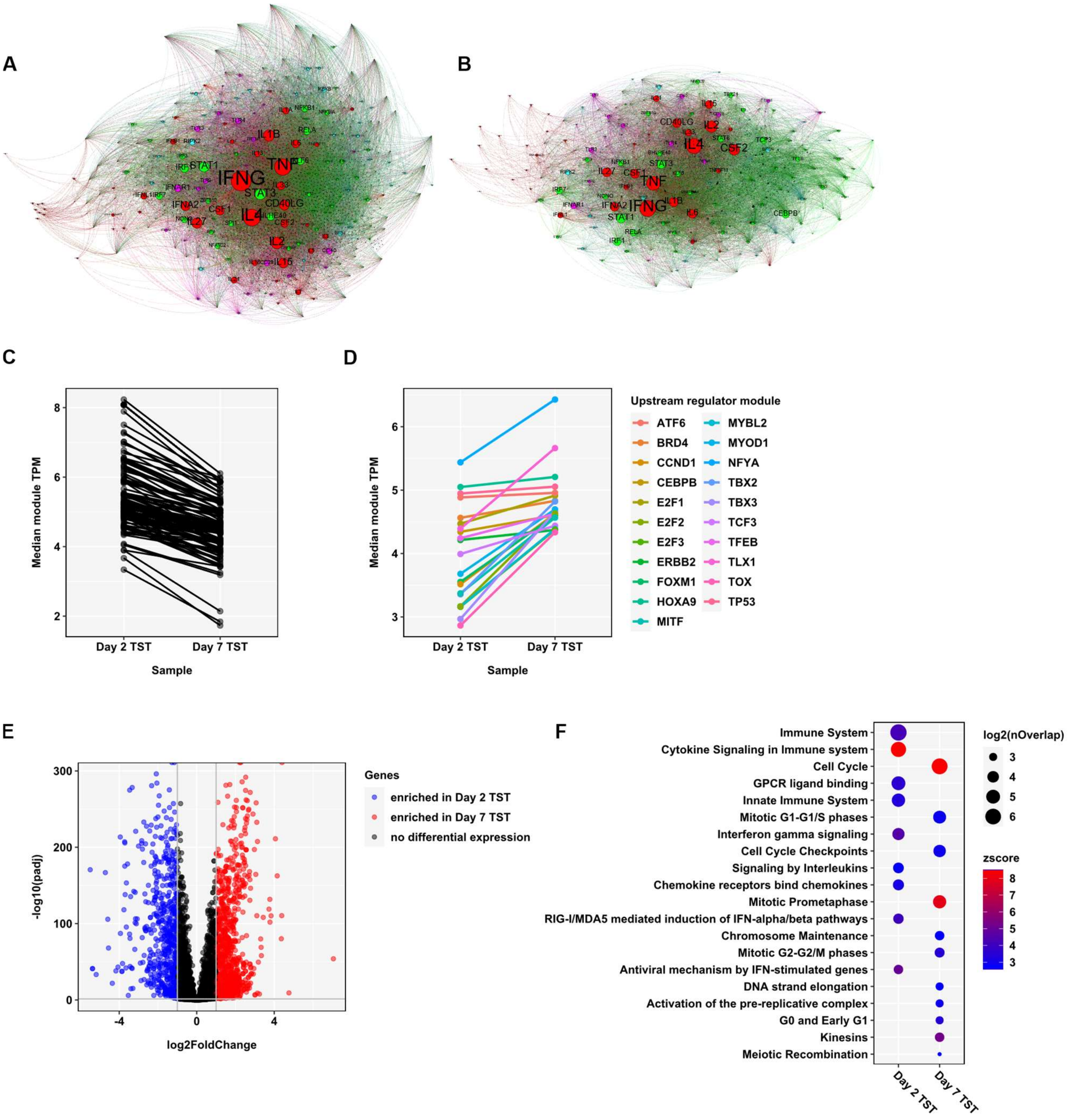

#### ***Increased cell cycle activity in Day 7 TSTs.***

Bulk RNA sequencing of n=33 saline controls, n=216 Day 2 TST and n=158 Day 7 TST samples was used to define the TST response. **(A-B)** Network diagrams of statistically significant regulators (FDR <0.05; labelled nodes), predicted to act upstream of genes (black nodes) enriched in either Day 2 TSTs (A) or Day 7 TSTs (B) compared to saline-injected control skin. Upstream regulators were stratified by molecular function (different colour of nodes), with node size proportional to -log10 p value. Nodes were clustered using Force Atlas 2 algorithm in Gephi (version 0.9.4). **(C-D)** Statistically significant upstream regulators (FDR <0.05) were identified for genes enriched in the comparison of all TST samples (Day 2 and Day 7) vs saline controls. The mean TPM (transcript per million) expression of the target genes for each upstream regulator (= module score) was quantified in individual Day 2 and Day 7 TST samples, and the median of these module scores calculated per group. Upstream regulators were stratified by whether the median score decreased **(C)** or increased **(D)** between Day 2 and Day 7. Each dot represents an upstream regulator module, showing the median score per group. In D, the legend lists the upstream regulator names, which are all associated with cell cycle activity. **(E)** Volcano plot showing statistical significance against quantitative gene expression differences between Day 2 and Day 7 TST samples, amongst 4150 integrated genes that are enriched in either Day 2 or Day 7 TSTs compared to saline controls. Genes highlighted in red and blue are upregulated on Day 2 (n=608) and Day 7 (n=1003), respectively, with an FDR <0.05 and fold change ≥2. **(F)** Enrichment of Reactome pathways among the genes differentially expressed between Day 2 and Day 7 TSTs, as identified in E. Node size indicates number of the differentially expressed genes that are associated with each pathway. Node colour represents the statistical enrichment Z score.

**Supplementary Figure 3**

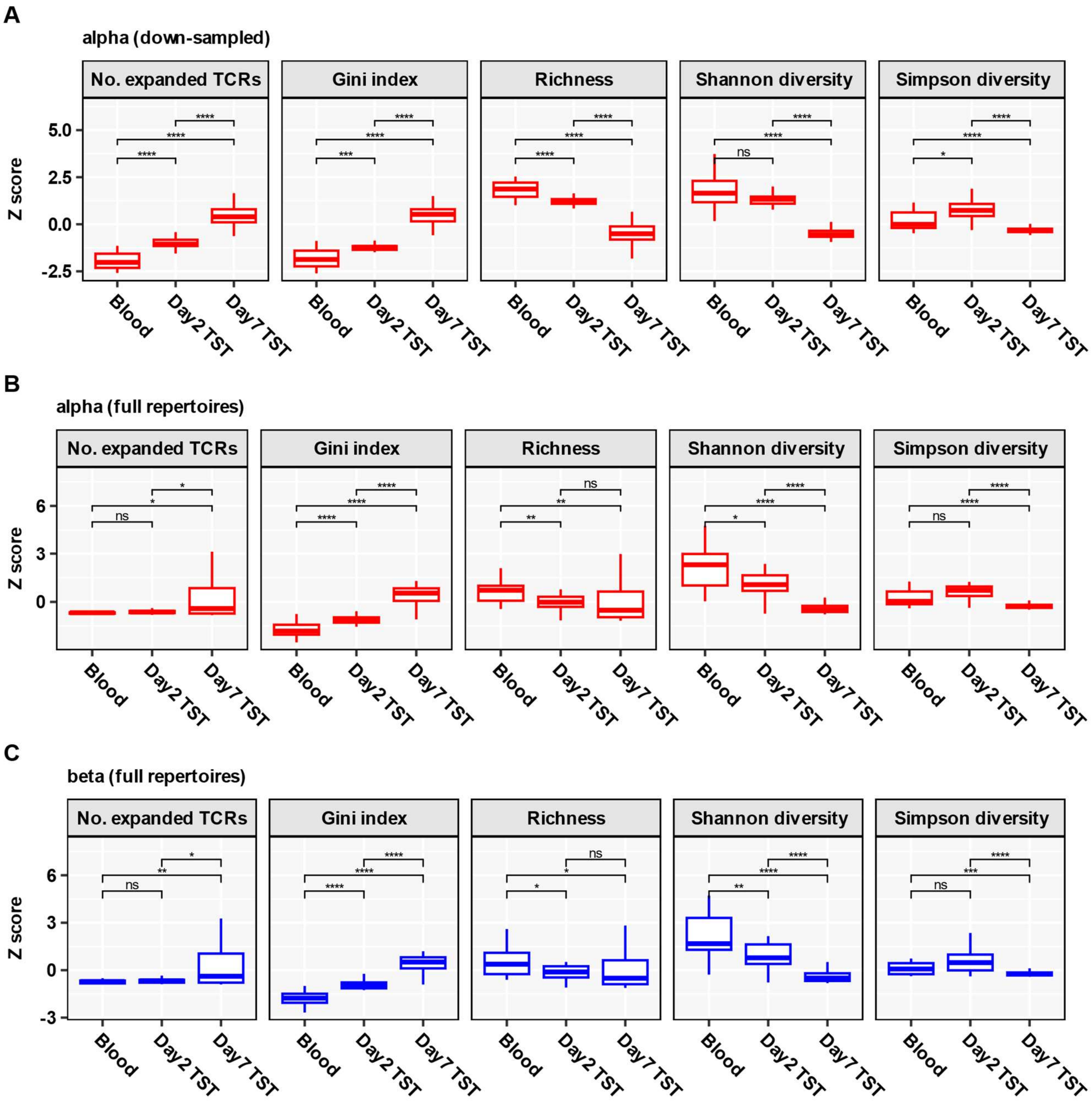

***TCR repertoire diversity metrics.***

Data are shown as Z-score values scaled across all samples, with boxplots depicting median and inter-quartile range (IQR), while outlier data points (more than 1.5*IQR beyond the box hinges) are shown as dots. Statistical significance was assessed with Wilcoxon tests and corrected for multiple testing (ns FDR>0.05, * FDR<0.05, ** FDR<0.01, *** FDR<0.01, **** FDR<0.0001). No. expanded TCRs = number of TCR sequences present more than once. **(A)** Diversity metrics calculated after individual alpha chain bulk TCR repertoires were down-sampled to 16,000 total TCRs (n=20 Blood, n=16 Day 2 TST, n=119 Day 7 TST). **(B-C)** Diversity metrics calculated in full repertoires (n=20 Blood, n=17 Day 2 TST, n=165 Day 7 TST), using either alpha **(B)** or beta **(C)** chain sequences.

.

**Supplementary Figure 4**

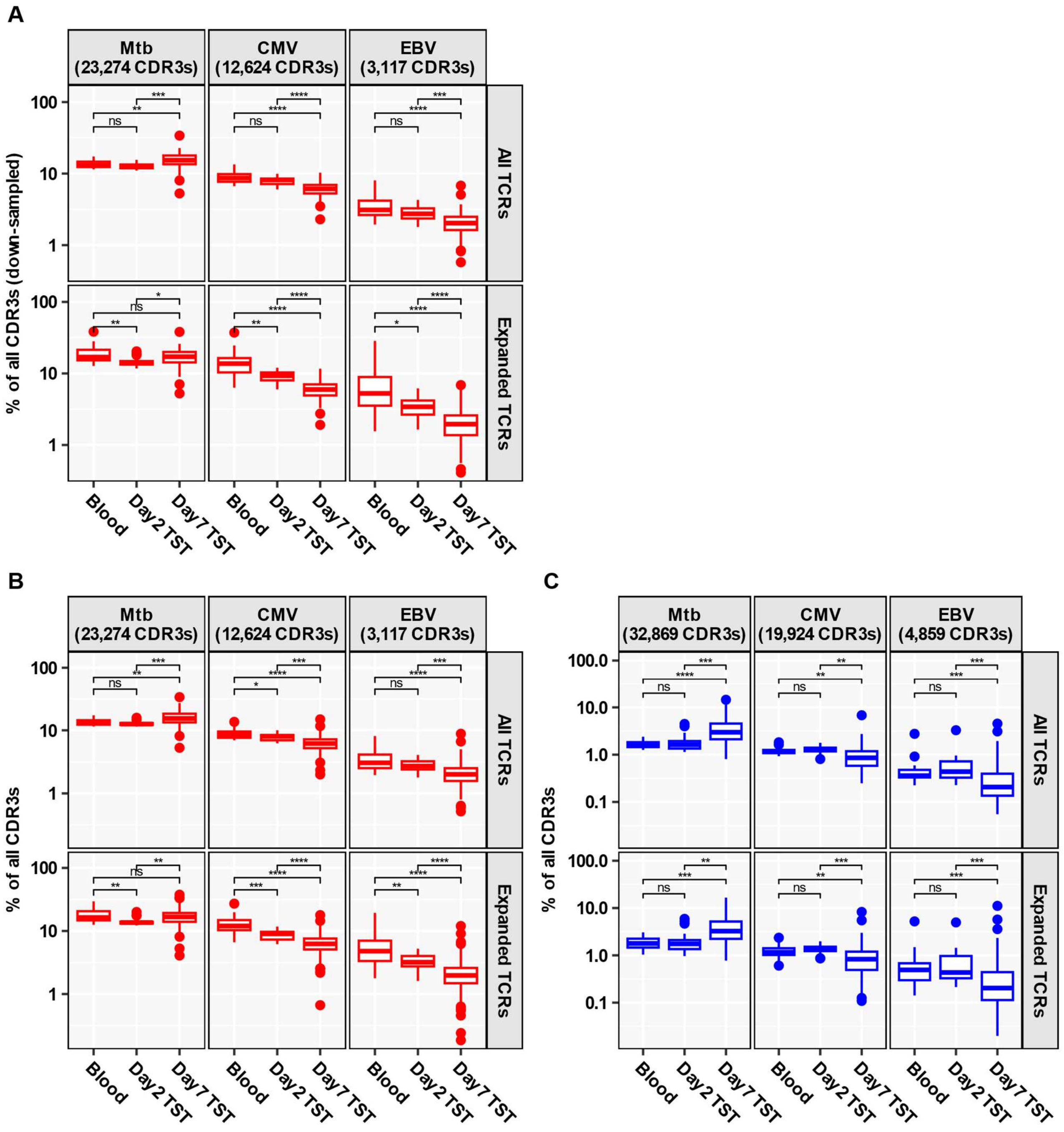

***Abundance of published antigen-reactive CDR3 sequences.***

Antigen-reactive CDR3s (specific for Mtb, CMV or EBV) were collated from VDJdb and McPAS databases as well as from Musvosvi et al., and their abundance in Blood and TST repertoires is shown as percentage of all TCRs or of all expanded TCRs (present more than once). The number of distinct published antigen-reactive CDR3s available to assess enrichment of antigen reactivity is indicated. Boxplots display median and inter-quartile range (IQR), with outlier data points (more than 1.5*IQR beyond the box hinges) shown as dots. Statistical significance was assessed with Wilcoxon tests and corrected for multiple testing (ns FDR>0.05, * FDR<0.05, ** FDR<0.01, *** FDR<0.01, **** FDR<0.0001). **(A)** Abundance calculated after individual alpha chain bulk TCR repertoires were down-sampled to 16,000 TCRs (n=20 Blood, n=16 Day 2 TST, n=119 Day 7 TST). **(B-C)** Abundance calculated in full repertoires (n=20 Blood, n=17 Day 2 TST, n=165 Day 7 TST), using either alpha **(B)** or beta **(C)** chain sequences.

**Supplementary Figure 5**

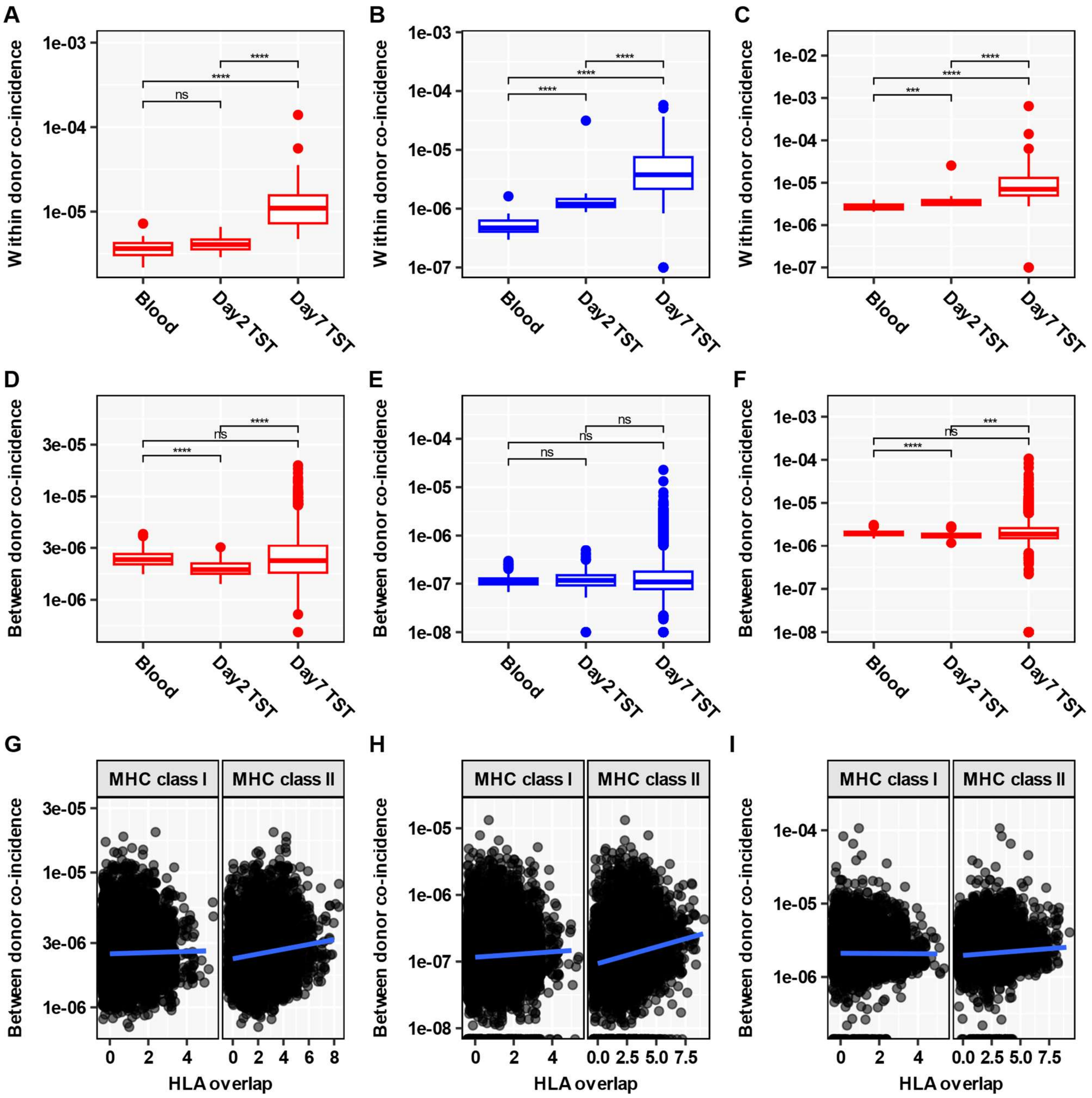

***Within- and cross-donor convergence of TCR repertoires.***

**(A-C)** Within-donor convergence of different TCR nucleotide sequences onto identical amino acid sequences in down-sampled alpha **(A)**, full beta **(B)** or full alpha **(C)** repertoires. The number of samples in down-sampled alpha repertoires **(A)** is n=20 Blood, n=16 Day 2 TST and n=119 Day 7 TST. The number of samples in full repertoires **(B-C)** is n=20 Blood, n=17 Day 2 TST and n=165 Day 7 TST. **(D-F)** Cross-donor TCR convergence, calculated in any two individuals, in down-sampled alpha **(D)**, full beta **(E)** or full alpha **(F)** repertoires. The number of pairwise comparisons for down-sampled alpha repertoires **(D)** is n=190 Blood, n=120 Day 2 TST and n=7021 Day 7 TST. The number of pairwise comparisons for full repertoires **(E-F)** is n=190 Blood, n=136 Day 2 TST and n=13530 Day 7 TST. **(G-I)** Cross-donor TCR convergence in Day 7 TSTs, stratified by the number of class 1 or class 2 HLA alleles shared between any two individuals, using down-sampled alpha **(G)**, full beta **(H)** or full alpha **(I)** repertoires.

The boxplots in A-F depict median and inter-quartile range (IQR) for each group, with outlier data points (more than 1.5*IQR beyond the box hinges) shown as dots. Statistical significance was assessed with Wilcoxon tests and corrected for multiple testing (ns FDR>0.05, *** FDR<0.001, **** FDR<0.0001). In G-I, each dot represents a pairwise comparison, with linear regression line shown in blue.

**Supplementary Figure 6**

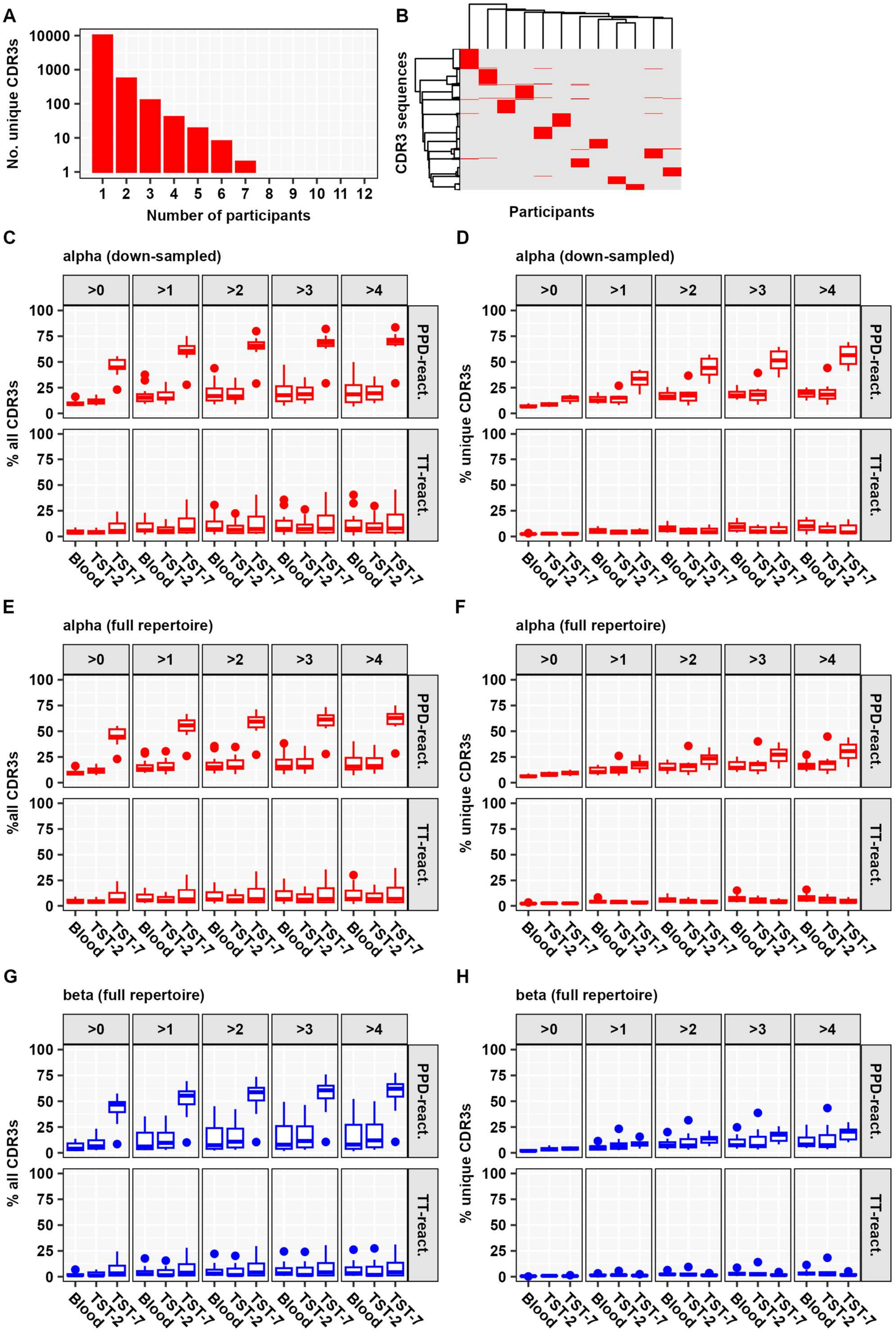

***Expansion of private Mtb-reactive TCRs in Day 7 TSTs.***

PBMCs from a subset of participants (n=12) were stimulated in vitro with one of 10 μg/ml purified protein derivative of Mtb (PPD), 100 μg/ml tetanus toxoid (TT), or control media for 6 days. Antigen-reactive CDR3s were identified by bulk TCR sequencing as CDR3s with ≥8-fold increase in count in PPD or TT cultures, but not unstimulated control cultures, compared to ex vivo blood from the same individual. **(A)** Histogram showing the number of unique in vitro PPD-reactive alpha chain CDR3s shared by different numbers of participants, demonstrating that the majority of PPD-reactive CDR3s are private (= found in only single participants). **(B)** Heatmap of unique in vitro PPD-reactive alpha chain CDR3s per participant. The dendrogram depicts Ward D2 linkage clustering. **(C-D)** Abundance of private PPD- or TT-reactive CDR3 sequences in down-sampled alpha chain repertoires from blood and TSTs, stratified by clone size (= TCR count). Abundance was quantified as percentage of total **(C)** or unique **(D)** CDR3s. **E-F.** Abundance of private PPD- or TT-reactive CDR3 sequences in full beta chain repertoires from blood and TSTs; quantified as percentage of total **(E)** or unique **(F)** CDR3s. **(G-H)** Abundance of private PPD- or TT-reactive CDR3 sequences in full alpha chain repertoires from blood and TSTs; quantified as percentage of total **(G)** or unique **(H)** CDR3s. The boxplots in C-H display median and inter-quartile range (IQR), with outlier data points (more then 1.5*IQR beyond the box hinges) shown as dots. Only participants used for paired in vitro stimulation experiments were used in this analysis (n=12 Blood, n=11 Day 2 TST, n=10 Day 7 TST).

**Supplementary Figure 7**

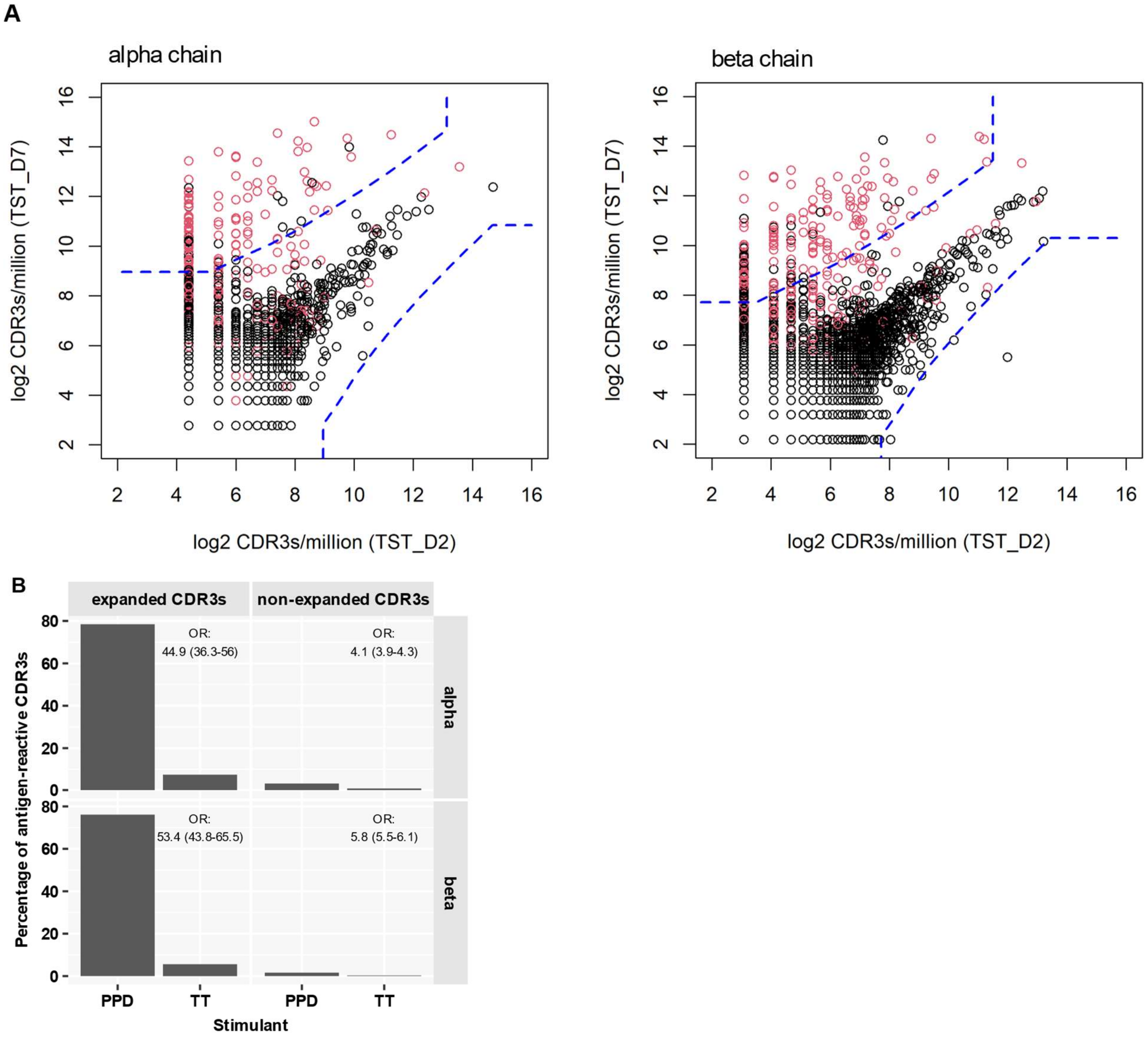

***Selective expansion of Mtb-reactive CDR3s between paired Day 2 and Day 7 TST samples.***

**(A)** A representative example of the pairwise comparison between Day 2 and Day 7 TST TCRs from the same individual. Each point is an individual CDR3 sequence, with its abundance on Day 7 plotted versus its abundance on Day 2. CDR3s absent in one of the paired samples were replaced with the median CDR3 count of that sample, and all CDR3 counts were then log-normalised (log2 counts/million). Dots in red represent PPD-reactive CDR3s, defined with in vitro stimulation experiments (see Figures 3 and S6). The dashed blue lines indicate the significance thresholds ($p < 0.0001$) for a Poisson distribution of counts, with CDR3s that fall to the left and above the dashed line considered expanded on Day 7. **(B)** Expanded and non-expanded CDR3s between paired Day 2 and Day 7 TSTs were integrated across n=10 individuals, stratified by chain, and their PPD and TT reactivity assessed using antigen-reactive CDR3s defined after in vitro stimulation (see Figures 3 and S6). Odds ratios (OR) with 95% confidence intervals were calculated to quantify enrichment of PPD vs TT reactivity amongst expanded and non-expanded clones.

**Supplementary Figure 8**

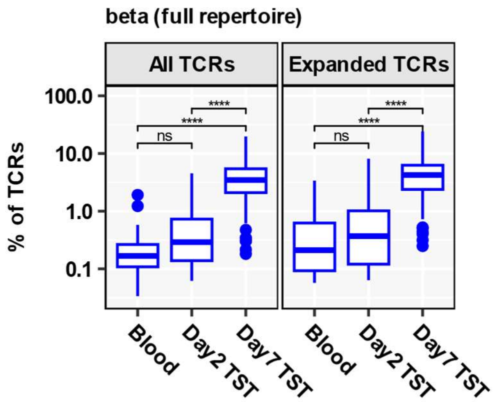

***Abundance and TB association of CDR3β metaclones in full repertoire of TCR sequences.***

In the full repertoire of TCRβ chain sequences, abundance of HLA class 2-restricted metaclones in individual blood or TST TCR repertoires, shown as percentage of all TCRs or of all expanded TCRs (present more than once). N=20 Blood, N=17 Day 2 TST, N=165 Day 7 TST. Boxplots display median and inter-quartile range (IQR), with outlier data points (more than 1.5*IQR beyond the box hinges) shown as dots. Statistical significance was assessed with Wilcoxon tests and corrected for multiple testing (ns FDR>0.05, **** FDR<0.0001).

**Supplementary Figure 9**

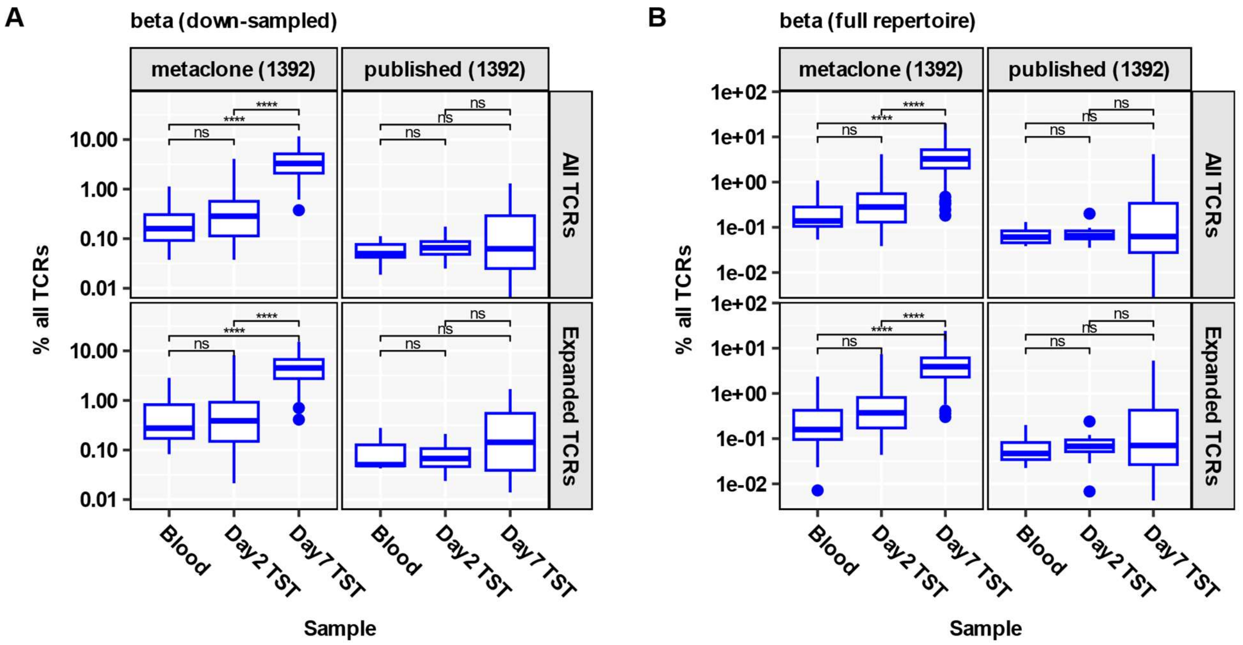

***Metaclones are more frequent in Day 7 TST compared to published Mtb-reactive CDR3 sequences.***

Abundance of HLA class 2 restricted metaclone CDR3s and equal numbers of randomly sampled published Mtb-reactive CDR3s in individual blood and TST repertoires, shown as percentage of all TCRs or of all expanded TCRs (present more than once). **(A)** Blood and TST repertoires were down-sampled to 16,000 $\beta$ chain TCRs each from N=20 Blood, N=16 Day 2 TST, N=128 Day 7 TST samples. **(B)** Full $\beta$ chain repertoires were used (N=20 Blood, N=17 Day 2 TST, N=165 Day 7 TST). Boxplots display median and inter-quartile range (IQR), with outlier data points (more than 1.5*IQR beyond the box hinges) shown as dots. Statistical significance was assessed with Wilcoxon tests and corrected for multiple testing (ns FDR>0.05, * FDR<0.05, **** FDR<0.0001).

**Supplementary Figure 10**

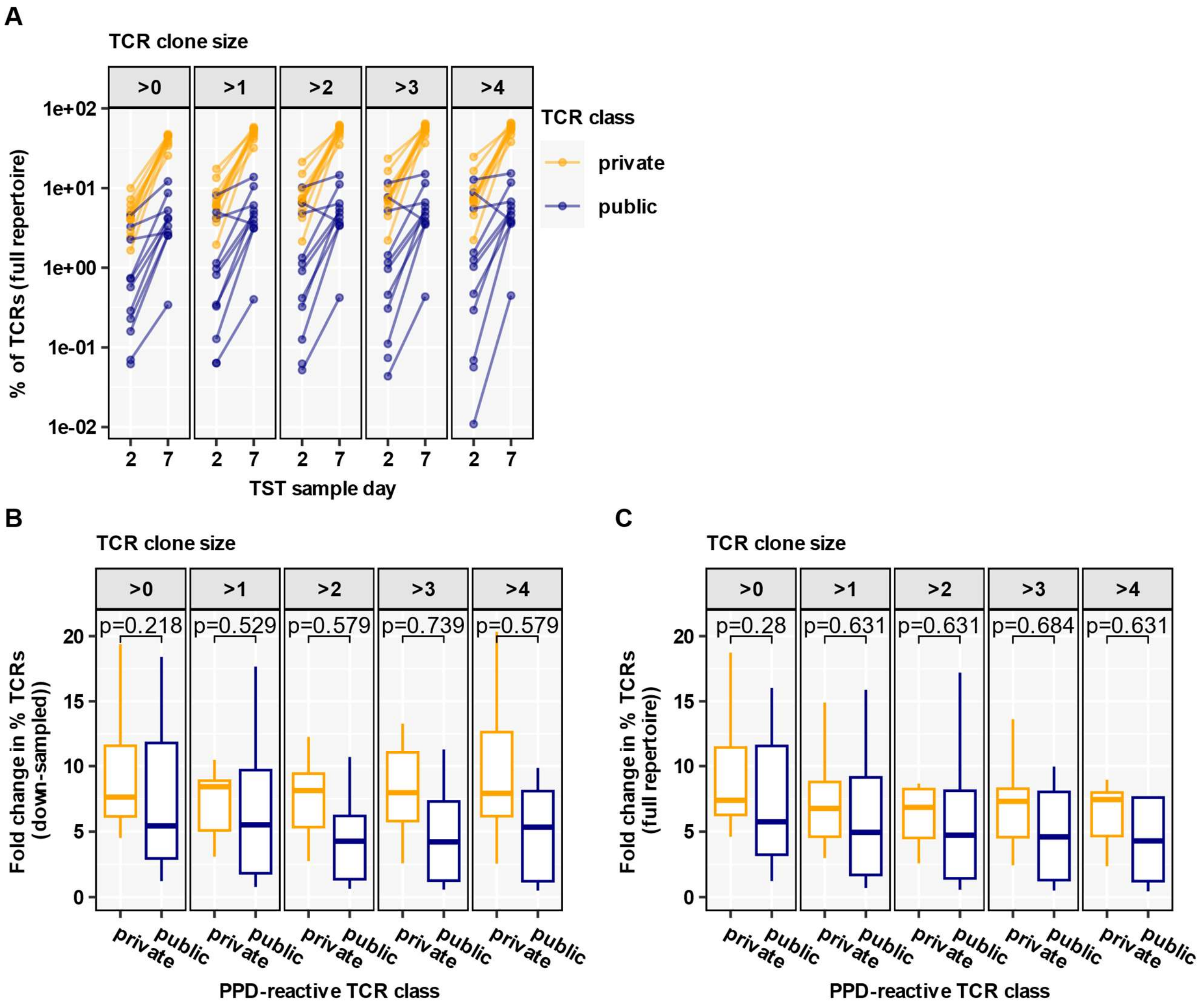

***Mtb-reactive metaclones constitute a small proportion of the Day 7 TST repertoire.***

**(A)** In the full repertoire of β chain TCR sequences, abundance of 'public' and 'private' Mtb-reactive CDR3s in the same individual was quantified as percentage of all Day 2 or all Day 7 TST TCRs, stratified by clone size (= TCR count). β–chain TCRs were classified as 'public' if they matched a Metaclonotypist metaclone, or else as 'private' if they matched a private PPD-reactive CDR3 sequence identified from ex vivo stimulated PBMC from the same individual (see Figure 3). CDR3s matching both a metaclone and private PPD-reactive CDR3 were classed as 'public'. This analysis was restricted to individuals with paired in vitro stimulation experiments (n=11 Day 2 TST, n=10 Day 7 TST). **(B-C)** Fold-change in private and public CDR3s in day 7 compared to day 2 TSTs stratified by clone size, using repertoires down-sampled to 16,000 TCRs each (B) or full repertoires (C). Box and whisker plots represent median, interquartile range and 1.5 IQR limits for N=10 individuals. P values shown for Wilcoxon tests for each analysis stratified clone size.

**Supplementary Figure 11**

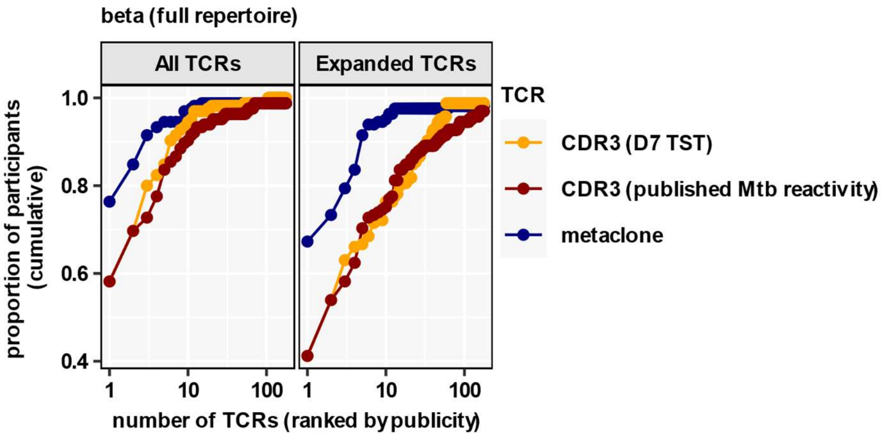

***Publicity of Mtb-reactive TCRs in the Day 7 TST.***

In the full repertoire of TCR$\beta$ chain sequences, HLA class 2-restricted metaclones (blue), CDR3s with published Mtb reactivity (red), and CDR3s present in Day 7 TSTs (yellow) were each ranked by their publicity across all available Day 7 TST repertoires (n=165) and plotted against the cumulative proportion of participants expressing the TCR. Presence of TCRs was assessed using either all TCR sequences in each sample or only expanded TCRs (present more than once).
